# Glycogen synthase kinase-3 is essential for T_reg_ development and function

**DOI:** 10.1101/2024.10.03.616474

**Authors:** Matheswaran Kandasamy, Hana F. Andrew, Iwan G. Raza, Robert Mitchell, Mariana Borsa, Moustafa Attar, Alexander J Clarke

**Affiliations:** Kennedy Institute of Rheumatology, University of Oxford, Roosevelt Drive, Oxford, OX3 7FY

## Abstract

T_regs_ are critical regulators of the immune response, but the cellular signalling pathways that control their development and homeostasis remain to be determined. We found that glycogen synthase kinase-3 (GSK3), a kinase which integrates signals from AKT and mTOR, was essential for T_reg_ development, restraining fatal autoimmunity. Loss of *Gsk3* led to metabolic rewiring in T_regs_, with disordered nucleotide metabolism and activation of OxPhos. Acute deletion of *Gsk3* did not affect T_reg_ frequency or numbers, but induced an effector gene expression program, and led to the formation of populations with pro-inflammatory signatures. The loss of *Gsk3* in T_regs_ profoundly enhanced anti-tumoral immune responses and suppressed tumour growth.

## Introduction

Regulatory T cells (T_reg_) expressing forkhead box P3 (FOXP3) are essential to control self-reactivity and maintain immune tolerance^1^. Whilst their function in restraining inflammation is clear, they may also suppress anti-tumoral immune responses and so promote tumour growth^2^. During their development, activation, and homeostasis, T_regs_ undergo profound alterations in their metabolic phenotype. Critical to this programming is signalling through the PI3K-AKT-mTOR axis. Phosphoinositide 3-kinase (PI3K) transduces signals from a variety of receptors, including the T cell receptor (TCR), and its key effector is AKT, phosphorylated by pyruvate dehydrogenase kinase-1 (PDK1) or mechanistic target of rapamycin complex 2 (mTORC2)^3,4^. Following phosphorylation, AKT then phosphorylates and inhibits tuberous sclerosis complex 2 (TSC2), to activate mTORC1 signalling^5,6^. Each component of PI3K-Akt-mTOR signalling modulates T_reg_ development and function^7–12^. However, knowledge of downstream effector pathways remains incomplete.

A critical interactor of the PI3K-AKT-mTOR pathway is glycogen synthase kinase-3 (GSK3)^13^. GSK3 is a serine/threonine kinase which exists in two paralogs, GSK3ɑ and GSK3β, which share a highly conserved catalytic region, but are encoded by separate genes. Initially characterised as phosphorylating glycogen synthase in response to insulin signalling, GSK3 has subsequently been identified as having a wide range of substrates with diverse roles in metabolism, cell signalling, and the cell cycle^14^. AKT phosphorylates serine 21 or serine 9 of GSK3ɑ and GSK3β respectively, inhibiting binding of their substrates and therefore kinase activity. AKT activity in T cells has been shown to inhibit T_reg_ formation, and T_regs_ themselves have lower levels of AKT phosphorylation than effector T cells^9,10^. GSK3 also phosphorylates phosphatase and tensin homolog (PTEN), reducing its function, which is required for T_reg_ suppressive capacity and lineage stability^11^. The mTORC2 component RPTOR independent companion of mTOR complex 2 (RICTOR) is a target of GSK3, and relative mTORC2 activity tunes mTORC1 to control T_reg_ function^8,11,15,16^.

A key substrate of GSK3 is β-catenin, which is degraded by the β-catenin destruction complex comprised of GSK3, AXIN, adenomatous polyposis coli (APC), and casein kinase 1ɑ (CK1ɑ). GSK3 phosphorylates AXIN, stabilising the complex and so regulating β-catenin degradation^17^. β-catenin has been shown to enhance the suppressive capacity of T_regs_ in some studies, but to be detrimental in others^18–22^.

In effector T cells, GSK3 is important for activation, and regulation of differentiation^23–27^. Notably, either pharmacological inhibition or deletion of GSK3 leads to increased tumour killing^24,28–30^. Pharmacologic GSK3 inhibition *in vitro* enhances FOXP3 expression and/or suppressive function in both murine and human T_regs_, in most but not all studies^22,31–35^. This suggests that GSK3 activity may be an intrinsic negative regulator of T_reg_ function.

Collectively these observations place GSK3 as a potentially key regulatory node in T_reg_ physiology. However, whether GSK3 is required for T_reg_ development, function, and homeostasis *in vivo* is unknown.

We found that GSK3 was essential for T_reg_ development, preventing fatal systemic autoimmunity. GSK3 is required to maintain expression of key suppressive molecules in T_regs_, and to regulate metabolic homeostasis, restraining OxPhos. Acute deletion of GSK3, whilst initiating upregulation of genes typically required for T_reg_ activity, led to defective function, and enhanced tumour killing. These findings highlight GSK3 as an essential regulator of T_reg_ function and provide a rationale for the use of GSK3 inhibition to enhance anti-tumoral immune responses.

## Results

### Deletion of Gsk3a/b in T_regs_ results in fatal autoimmune disease

To determine the role of GSK3 in T_regs_, we generated *Foxp3*^YFP-Cre^ × *Gsk3a*^LoxP^ × *Gsk3b*^LoxP^ mice (hereafter Gsk3a/b^ΔTreg^ mice), in which both paralogs were conditionally deleted in T_regs_ using Foxp3^YFP-Cre^, which could also be identified by their expression of YFP (Extended Data Fig. 1A).

Unexpectedly, Gsk3a/b^ΔTreg^ mice succumbed at around three weeks following birth, from a severe multisystem inflammatory disease affecting skin, lungs, and liver (Fig. 1A). Mice lacking only a single paralog (*Foxp3*^YFP-Cre^ × *Gsk3a*^LoxP^ or *Gsk3b*^LoxP^) were protected and developed normally, as did *Foxp3*^YFP-Cre^ mice heterozygous for *Gsk3a*^LoxP^ and *Gsk3b*^LoxP^ (which were thereafter used as controls). Gsk3a/b^ΔTreg^ mice grew poorly, developed dermatitis of the tail, and had dense inflammatory infiltrates in the liver and lungs, with lymph node and splenic enlargement (Fig. 1B, Extended Data Fig. 1B-D). Serum cytokine profiling revealed increased levels of IFN-γ, TNF-ɑ, MCP-1, IL-6, and IL-23 (Fig. 1C). Examination of spleen, mesenteric lymph node (MLN) and lungs by flow cytometry demonstrated accumulation of activated (CD44^hi^CD62L^lo^) T_conv_ cells, which produced high levels of IFN-γ and TNF-ɑ, and in the lungs IL-17A, following stimulation with PMA and ionomycin (Fig. 1D-J). IL-4 was not detected. Gsk3a/b^ΔTreg^ mice had evidence of humoral autoimmunity, with increased total serum IgM, IgG2a, IgG2b, and IgG2a and IgG2b anti-double-stranded DNA autoantibodies (Fig. 1K-M and Extended Data Fig. 1E). These were associated with increased germinal centre (GC) B cell and T follicular helper cell (T_FH_) numbers in mesenteric and inguinal lymph nodes (Fig. 1N-P and Extended Data Fig 1F-H). Mice lacking both GSK3 paralogs in T_regs_ therefore develop severe spontaneous autoimmunity associated with Th1, Th17, and humoral immune responses.

**Figure 1.**
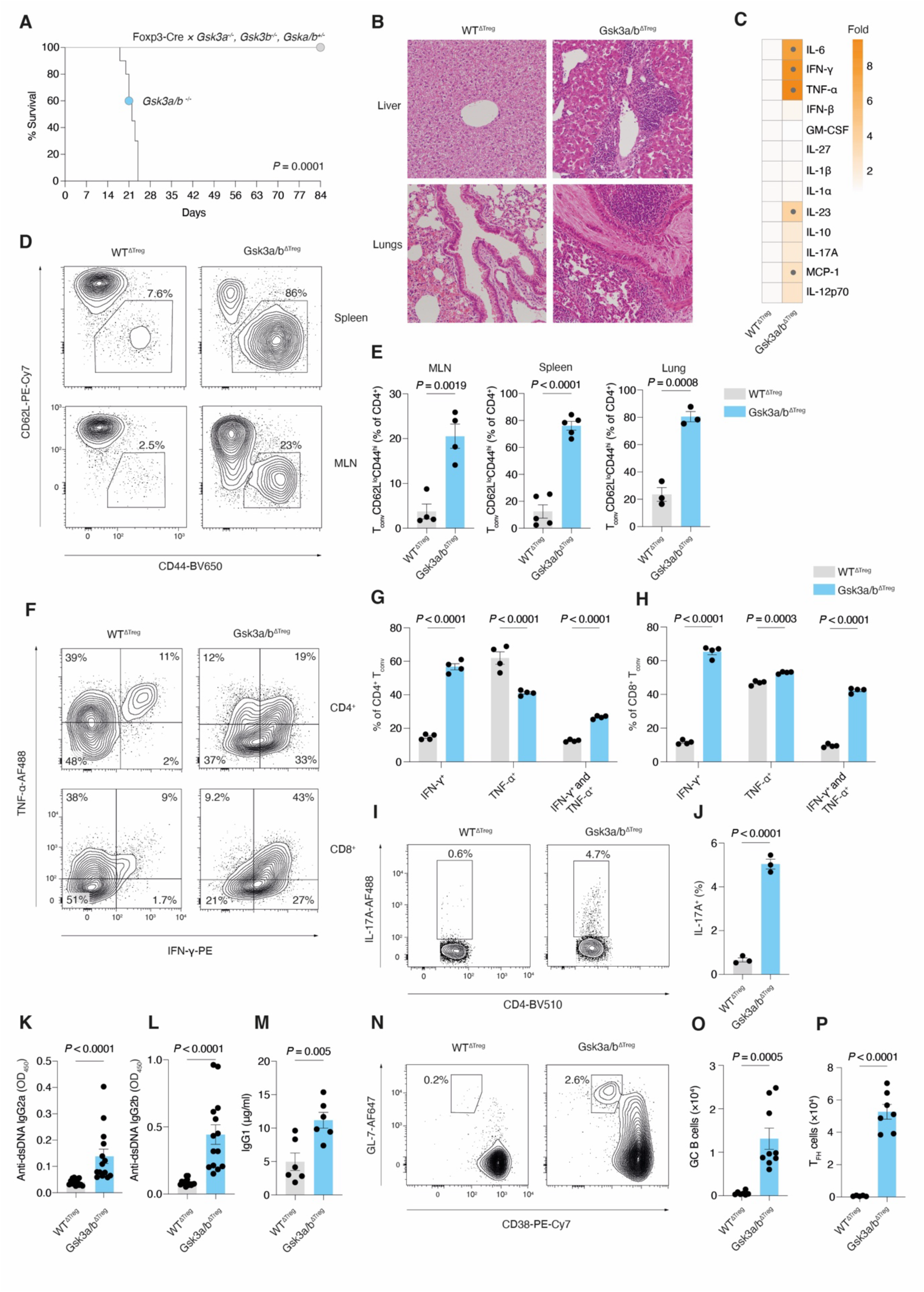
Loss of GSK3a/b in regulatory T cells induces fatal autoimmune disease. **A.** Survival curve for mice of the indicated genotype (*Foxp3-Cre*^+/-^ × Gsk3a^LoxP/+^ Gsk3b^LoxP/LoxP^ [n=4], *Foxp3-Cre*^+/-^ × Gsk3a^LoxP/LoxP^Gsk3b^LoxP/+^ [n=4], *Foxp3-Cre*^+/-^ × Gsk3a^LoxP/+^Gsk3b^LoxP+^ [n=4], and *Foxp3-Cre*^+/-^ × Gsk3a^LoxP/LoxP^Gsk3b^LoxP/LoxP^ [n=10]). **B.** Representative histology of liver and lungs of WT^ΔTreg^ and Gsk3a/b^ΔTreg^ mice at 3 weeks. **C.** Heatmap of relative fold change of indicated serum cytokines in Gsk3a/b^ΔTreg^ compared with WT^ΔTreg^ mice. Grey dots indicate *Padj* < 0.05 (n=8 mice per group). Pooled from >5 independent experiments. **D.** Representative flow cytometry plots of CD4^+^CD62L^lo^CD44^hi^ activated T cells from spleen and mesenteric lymph nodes (MLN) of WT^ΔTreg^ and Gsk3a/b^ΔTreg^ mice. **E.** Quantification of CD4^+^CD62L^lo^CD44^hi^ activated T cells from MLN, spleen, and lungs of WT^ΔTreg^ and Gsk3a/b^ΔTreg^ mice (n=3-5 per group). Representative of two independent experiments). **F.** Representative flow cytometry plots of splenic interferon-γ (IFN-γ^+^) and TNF-ɑ^+^ CD4^+^ and CD8^+^ T cells in WT^ΔTreg^ and Gsk3a/b^ΔTreg^ mice. **G.** Quantification of splenic CD4^+^ IFN-γ^+^ and TNF-ɑ^+^ T cells in WT^ΔTreg^ and Gsk3a/b^ΔTreg^ mice as in **F** (n=4 per group). Representative of 3 independent experiments with similar results. **H.** Quantification of splenic CD8^+^ IFN-γ^+^ and TNF-ɑ^+^ T cells in WT^ΔTreg^ and Gsk3a/b^ΔTreg^ mice as in **F** (n=4 per group). Representative of 3 independent experiments. **I.** Representative flow cytometry plots of CD4^+^IL-17A^+^ T cells from lungs of WT^ΔTreg^ and Gsk3a/b^ΔTreg^ mice. **J.** Quantification of CD4^+^IL-17A^+^ T cells as in I (n=3 per group). Representative of two independent experiments. **K.** Quantification of IgG2a anti-dsDNA autoantibodies in sera of WT^ΔTreg^ and Gsk3a/b^ΔTreg^ mice by ELISA (n=14 per group). Pooled from >5 independent experiments. **L.** Quantification of IgG2b anti-dsDNA autoantibodies in sera of WT^ΔTreg^ and Gsk3a/b^ΔTreg^ mice by ELISA (n=14 per group). Pooled from >5 independent experiments. **M.** Quantification of serum IgG1 in sera of WT^ΔTreg^ and Gsk3a/b^ΔTreg^ mice by ELISA (n=14 per group). Pooled from >3 independent experiments. **N.** Representative flow cytometry plots of CD19^+^GL-7^+^CD38^-^ GC B cells from WT^ΔTreg^ and Gsk3a/b^ΔTreg^ MLN. **O.** Quantification of GC B cells from WT^ΔTreg^ and Gsk3a/b^ΔTreg^ MLN (n=7 per group. Pooled from 3 independent experiments. **P.** Quantification of CD4+PD-1^hi^CXCR5^hi^ T_FH_ cells from WT^ΔTreg^ and Gsk3a/b^ΔTreg^ MLN (n=7 per group). Pooled from 3 independent experiments. Statistical significance was determined by Mantel-Cox test (A), two-way ANOVA with Šidák’s multiple testing correction (C, G-H, K), or unpaired two-tailed t test (E, J, K-M, O-P). Data are presented as the mean +/- SEM. Each point represents a single mouse.

### GSK3 is required for T_reg_ development and homeostasis

We next examined T_reg_ populations in 3-week-old Gsk3a/b^ΔTreg^ mice (Fig. 2A). There was a drastic reduction in T_regs_ in both lymphoid tissue and thymus, with the exception of inguinal lymph nodes (Fig. 2B).

**Figure 2.**
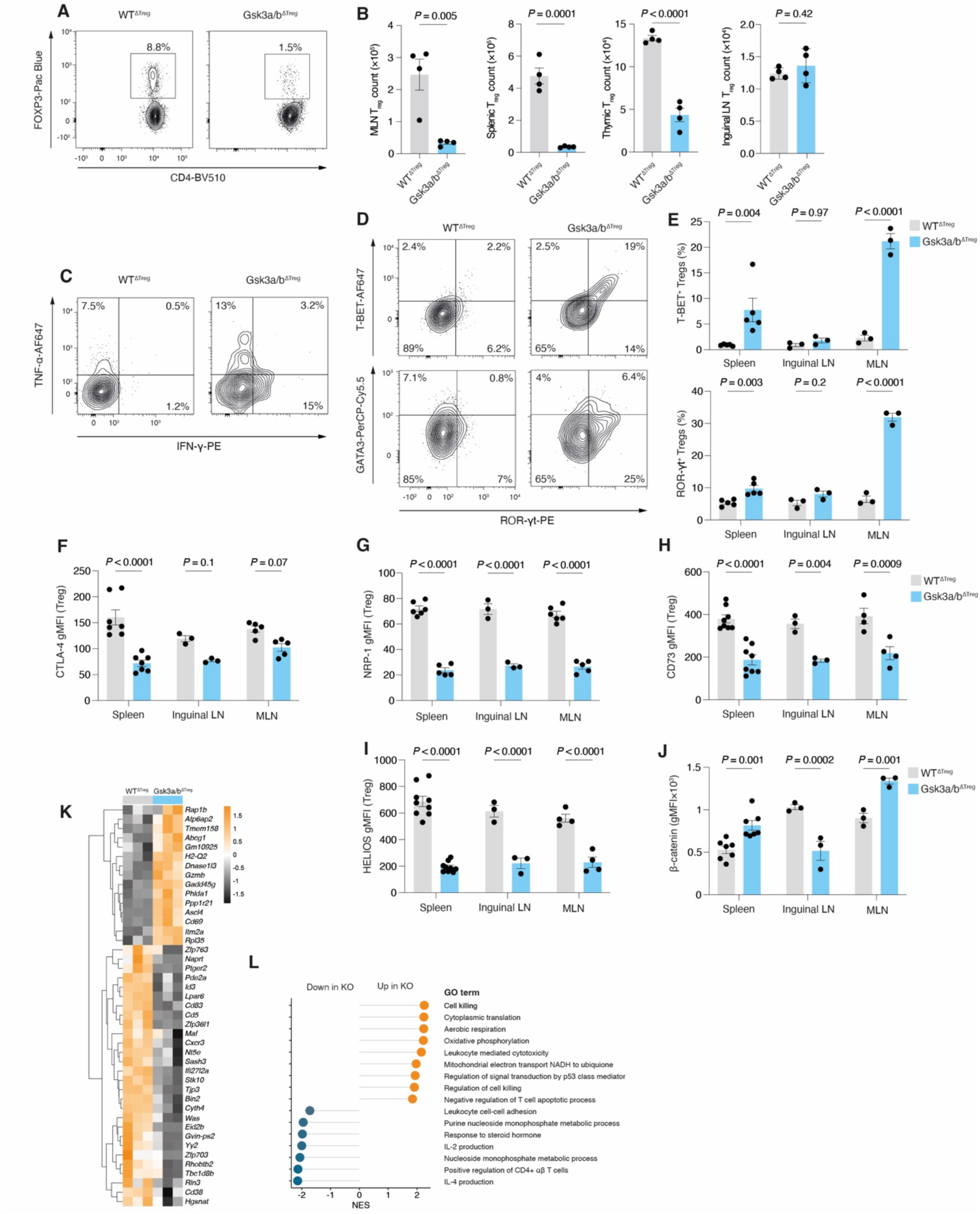
GSK3 is required for T_reg_ development and homeostasis. **A.** Representative flow cytometry plots of splenic TCRβ^+^CD4^+^FOXP3^+^ T_regs_ from WT^ΔTreg^ and Gsk3a/b^ΔTreg^ mice. **B.** Quantification of T_reg_ numbers from MLN, spleen, thymus, and inguinal LN of WT^ΔTreg^ and Gsk3a/b^ΔTreg^ mice (n=4 per group). Representative of 2 independent experiments. **C.** Representative flow cytometry plots of splenic IFN-γ^+^ and TNF-ɑ^+^ T_regs_ from WT^ΔTreg^ and Gsk3a/b^ΔTreg^ mice. **D.** Representative flow cytometry plots of splenic T-BET^+^, ROR γt^+^, and GATA3^+^ T_regs_ from WT^ΔTreg^ and Gsk3a/b^ΔTreg^ mice. **E.** Quantification of T-BET^+^ and ROR-γt T_regs_ in spleen, inguinal LN, and MLN from WT^ΔTreg^ and Gsk3a/b^ΔTreg^ mice (n=3-5 per group). Representative of 2 independent experiments. **F.** Quantification of CTLA-4 expression from T_regs_ in spleen, inguinal LN, and MLN from WT^ΔTreg^ and Gsk3a/b^ΔTreg^ mice (n=3-6 per group). Representative of 2 independent experiments. **G.** Quantification of NRP-1 as in F (n=5-6 per group). Representative of 2 independent experiments. **H.** Quantification of CD73 as in F (n=3-8 per group). Representative of 2 independent experiments. **I.** Quantification of HELIOS as in F (n=3-8 per group). Representative of 2 independent experiments. **J.** Quantification of β-catenin as in F (n=3-7 per group). Representative of 2 independent experiments. **K.** Heatmap of gene expression by RNA sequencing in splenic CD4^+^Foxp3-eGFP^+^ T_regs_ from WT^ΔTreg^ and Gsk3a/b^ΔTreg^ mice (n=3 per group). **L.** Gene set enrichment analysis (GSEA) of samples as in K. Normalised enrichment score for gene ontogeny (GO) terms shown. All pathways shown are statistically significant (*P*_adj_ <0.05). Statistical significance was determined by unpaired two-tailed t test (B) or two-way ANOVA with Šidák’s multiple testing correction (E -J). Data are presented as the mean +/- SEM. Each point represents a single mouse.

T_regs_ from Gsk3a/b^ΔTreg^ mice produced high levels of IFN-γ and TNF-ɑ, and expressed elevated levels of the transcription factors T-BET and ROR-γt, which control Th1 and Th17 programs respectively (Fig. 2C-E, Extended Data Fig. 2A). However, despite ROR-γt being increased, we did not detect significant IL-17A production in Gsk3a/b^ΔTreg^ T_regs_. The Th2 transcription factor GATA-3 was not upregulated, in keeping with its likely suppression by elevation of the transcription factor IRF4, (Extended Data Fig. 2B)^36^.

T_regs_ from Gsk3a/b^ΔTreg^ mice had decreased expression of surface molecules associated with suppressive capacity, including CTLA-4, NRP-1, CD73, and CD25 (Fig. 2F-H, Extended Data Fig. 2D-E). Also decreased were levels of the transcription factors HELIOS and FOXP3 which help maintain T_reg_ identify and stability (Fig. 2I and Extended Data Fig. 2C-D). As expected given the role of GSK3 in its degradation, β-catenin was increased in Gsk3a/b^ΔTreg^ T_regs_ (Fig. 2J and Extended Data Fig. 2C). CD25 (IL-2Rɑ chain) expression was lower in Gsk3a/b^ΔTreg^ T_regs_, and this was associated with impaired STAT-5 phosphorylation following stimulation with IL-2, which has been shown to compromise T_reg_ stability (Extended Data Fig. 2C-E)^37^.

Loss of *Gsk3* affected T_regs_ in a tissue site specific manner, with features of activation (CD44^hi^CD62L^lo^ phenotype) increased in inguinal lymph node, decreased in MLN, but with no difference in the spleen (Extended Data Fig. 2F).

We next sorted T_regs_ from the spleens of Gsk3a/b^ΔTreg^ and WT^ΔTreg^ mice, and performed bulk RNA sequencing. We found differential expression of 99 genes, with upregulation of genes associated with effector T_reg_ function, Th1 polarisation, and tumour killing, notably *Gzmb* and *Gadd45g*^38^, and downregulation of suppressive genes including *Nt5e* which encodes 5’-nucleotidase (CD73), *Cd83*, *Cd38*, *Maf*, *Id3*, Cd5, and *Pde2a* (Fig. 2K). Other downregulated genes encoding regulatory proteins included *Cxcr3*, which negatively regulates T_reg_ function in the tumour microenvironment, and *Ptger2*, which encodes the prostaglandin E2 receptor.

Gene set enrichment analysis (GSEA) showed upregulation of cell killing pathways, and downregulation of IL-4 and IL-2 production pathways, and leukocyte cell-cell adhesion (Fig. 2L). Notably, there was strong upregulation of OxPhos and downregulation of nucleoside metabolic processes.

Deletion of Gsk3 therefore induces severe T_reg_ dysfunction, associated with loss of lineage stability and transcriptional features of altered metabolism.

### *Gsk3a/b* deletion leads to cell intrinsic effects in T_regs_

An inflammatory environment may lead to T_reg_ destabilisation, and given our finding of a severe numerical reduction in T_reg_ numbers in Gsk3a/b^ΔTreg^ mice, associated with a Scurfy-like phenotype, it remained unclear whether the defects we observed were cell intrinsic. To address this, we first generated T_regs_ from Gsk3a/b^ΔTreg^ and WT^ΔTreg^ CD4^+^ T cells *in vitro*, by culturing naïve CD4^+^ T cells in the presence of IL-2 and TGF-β for 6 days. There was a slight but non-statistically significant reduction in FOXP3^+^ T cells, and examination of *in vitro* T_reg_ surface markers showed concordance with the phenotype of *ex vivo* Gsk3a/b^ΔTreg^ T_regs_, with reduced expression of CD73 and NRP-1, and increased β-catenin (Extended Data Fig 3A-E). CTLA-4 expression was unchanged however. We found that *in vitro*-generated T_regs_ from Gsk3a/b^ΔTreg^ mice had markedly less capacity to suppress proliferation of wild type CD4^+^ T responder cells stimulated with anti-CD3 and CD28 in a co-culture assay, compared with those from WT^ΔTreg^ mice (Fig. 3A-B).

**Figure 3.**
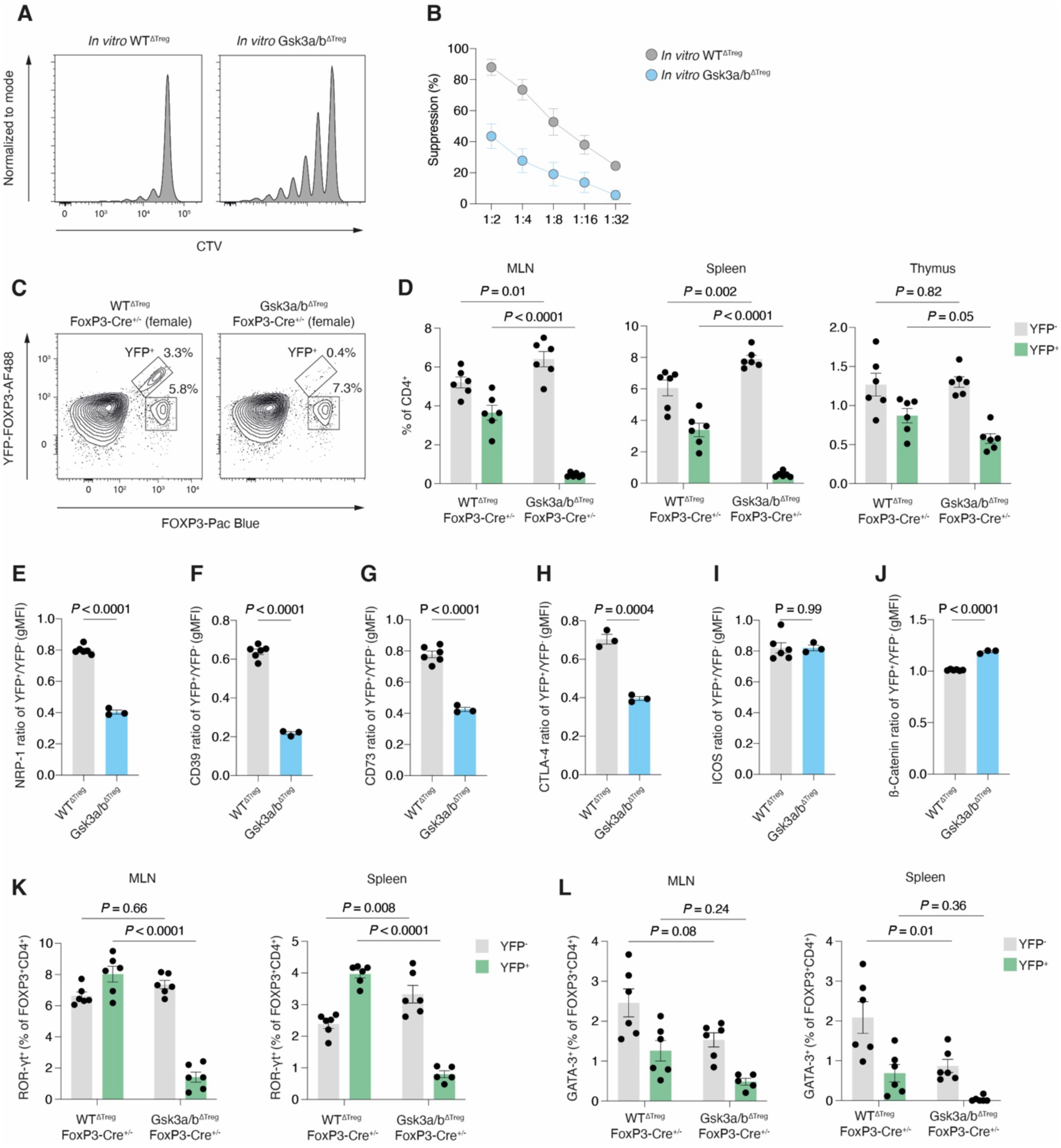
*Gsk3a/b* deletion leads to cell intrinsic effects in T_regs_. **A.** Representative histograms showing CellTrace Violet (CTV) dilution in divided responder T cells, cultured at a 4:1 responder:T_reg_ ratio with *in vitro*-generated T_regs_. **B.** Quantification of suppressive capacity of *in vitro*-generated T_regs_ from WT^ΔTreg^ and Gsk3a/b^ΔTreg^ mice, mixed with wild type responder T cells at the indicated ratio (n=4 per group). Representative of two independent experiments. **C.** Representative flow cytometry plots indicating gating strategy for mosaic T_regs_. Within CD4^+^TCRβ^+^ cells from female *Foxp3^YFP-Cre^* heterozygote × *Gsk3a/b^LoxP^* mice, YFP^+^FOXP3^+^ T_regs_ were considered to be *Gsk3a/b^-/-^*, whilst YFP^-^FOXP3^+^ were wild type. **D.** Quantification of frequencies of *Gsk3a/b^-/-^* and wild type mosaic T_regs_ in spleen, MLN, and thymus as in C (n=6 per group). Representative of two independent experiments. **E.** Ratio of gMFI of NRP-1 between *Gsk3a/b^-/-^* and wild type mosaic T_regs_ within individual mice (n=3 in *Gsk3a/b^LoxP^* × *Foxp3^YFP^-*Cre^+/-^ and n=6 in WT*^LoxP^* × *Foxp3^YFP^-*Cre^+/-^ mice). Representative of two independent experiments. **F.** Ratio of gMFI of CD39 between *Gsk3a/b^-/-^* and wild type mosaic T_regs_ within individual mice (n=3 in *Gsk3a/b^LoxP^* × *Foxp3^YFP^-*Cre^+/-^ and n=6 in WT*^LoxP^* × *Foxp3^YFP^-*Cre^+/-^ mice). Representative of two independent experiments. **G.** Ratio of gMFI of CD73 between *Gsk3a/b^-/-^* and wild type mosaic T_regs_ within individual mice (n=3 in *Gsk3a/b^LoxP^* × *Foxp3^YFP^-*Cre^+/-^ and n=6 in WT*^LoxP^* × *Foxp3^YFP^-*Cre^+/-^ mice). Representative of two independent experiments. **H.** Ratio of gMFI of CTLA-4 between *Gsk3a/b^-/-^* and wild type mosaic T_regs_ within individual mice (n=3 in *Gsk3a/b^LoxP^* × *Foxp3^YFP^-*Cre^+/-^ and n=3 in WT*^LoxP^* × *Foxp3^YFP^-*Cre^+/-^ mice). Representative o two independent experiments. **I.** Ratio of gMFI of ICOS between *Gsk3a/b^-/-^* and wild type mosaic T_regs_ within individual mice (n=3 in *Gsk3a/b^LoxP^* × *Foxp3^YFP^-*Cre^+/-^ and n=6 in WT*^LoxP^* × *Foxp3^YFP^-*Cre^+/-^ mice). Representative of two independent experiments. **J.** Ratio of gMFI of β-catenin between *Gsk3a/b^-/-^* and wild type mosaic T_regs_ within individual mice (n=3 in *Gsk3a/b^LoxP^* × *Foxp3^YFP^-*Cre^+/-^ and n=6 in WT*^LoxP^* × *Foxp3^YFP^-*Cre^+/-^ mice). Representative of two independent experiments. **K.** Quantification of ROR-γt^+^ population in *Gsk3a/b^-/-^* and wild type mosaic T_regs_ in MLN and spleen (n=6 in *Gsk3a/b^LoxP^* × *Foxp3^YFP^-*Cre^+/-^ and n=6 in WT*^LoxP^* × *Foxp3^YFP^-*Cre^+/-^ mice. Representative of two independent experiments. **L.** Quantification of GATA-3^+^ population in *Gsk3a/b^-/-^* and wild type mosaic T_regs_ in MLN and spleen (n=6 in *Gsk3a/b^LoxP^* × *Foxp3^YFP^-*Cre^+/-^ and n=6 in WT*^LoxP^* × *Foxp3^YFP^-*Cre^+/-^ mice. Representative of two independent experiments. Statistical significance was determined by two-way ANOVA with Šidák’s multiple testing correction (D, K-L) or unpaired two-tailed t test (E). Data are presented as the mean +/- SEM. Each point represents a single mouse.

Since *Foxp3* is encoded on the X chromosome, which is subject to random inactivation in female cells, natural mosaicism present in mice heterozygous for *Foxp3^YFP^-*Cre can be used to examine the effects of *Gsk3a/b* deletion in a chimeric setting, in which T_regs_ are sufficiently abundant and in which cell intrinsic effects can be observed in an undisturbed environment. We found that YFP^+^ T_regs_ from female *Gsk3a/b^LoxP^* × *Foxp3^YFP^-*Cre^+/-^ mice were strikingly underrepresented proportionately and numerically in the FOXP3^+^ population from spleen and MLN, and to a lesser extent the thymus (Fig. 3C-D and Extended Data Fig. 3F). They also had a more naïve phenotype, with a lower proportion of CD62L^lo^CD44^hi^ T_regs_(Extended Data Fig. 3G)

The expression of suppressive surface proteins in YFP^+^ mosaic T_regs_ largely mirrored Gsk3a/b^ΔTreg^ mice, with reduction in NRP-1, CD39, CD73, CTLA-4, but not ICOS, and with an increase in β-catenin (Fig. 3E-J). Interestingly, and in contrast to Gsk3a/b^ΔTreg^ T_regs_, there was a large reduction in the expression of ROR-γt in both MLN and splenic YFP^+^ mosaic T_regs_, seen to a lesser extent with GATA-3 (Fig. 3K-L). We did not detect T-BET expression, presumably due to lack of systemic inflammation.

The defects seen in T_reg_ homeostasis following *Gsk3a/b* deletion are thus in major part cell intrinsic, but also reflect a failure to suppress more widespread Th1-driven immunity once inflammation is initiated.

### Gsk3a/b deletion rewires T_reg_ metabolism

GSK3 is an important node in control of metabolism, and our transcriptional profiling demonstrated a significant enrichment of pathways related to OxPhos and aerobic respiration, and downregulation of nucleotide metabolic pathways in Gsk3a/b^ΔTreg^ T_regs_. We first quantified proteins of the electron transport chain (ETC) in splenic T_regs_. There was upregulation of cytochrome c oxidase subunit IV (COX IV) and cytochrome *c* (Fig. 4A).

**Figure 4.**
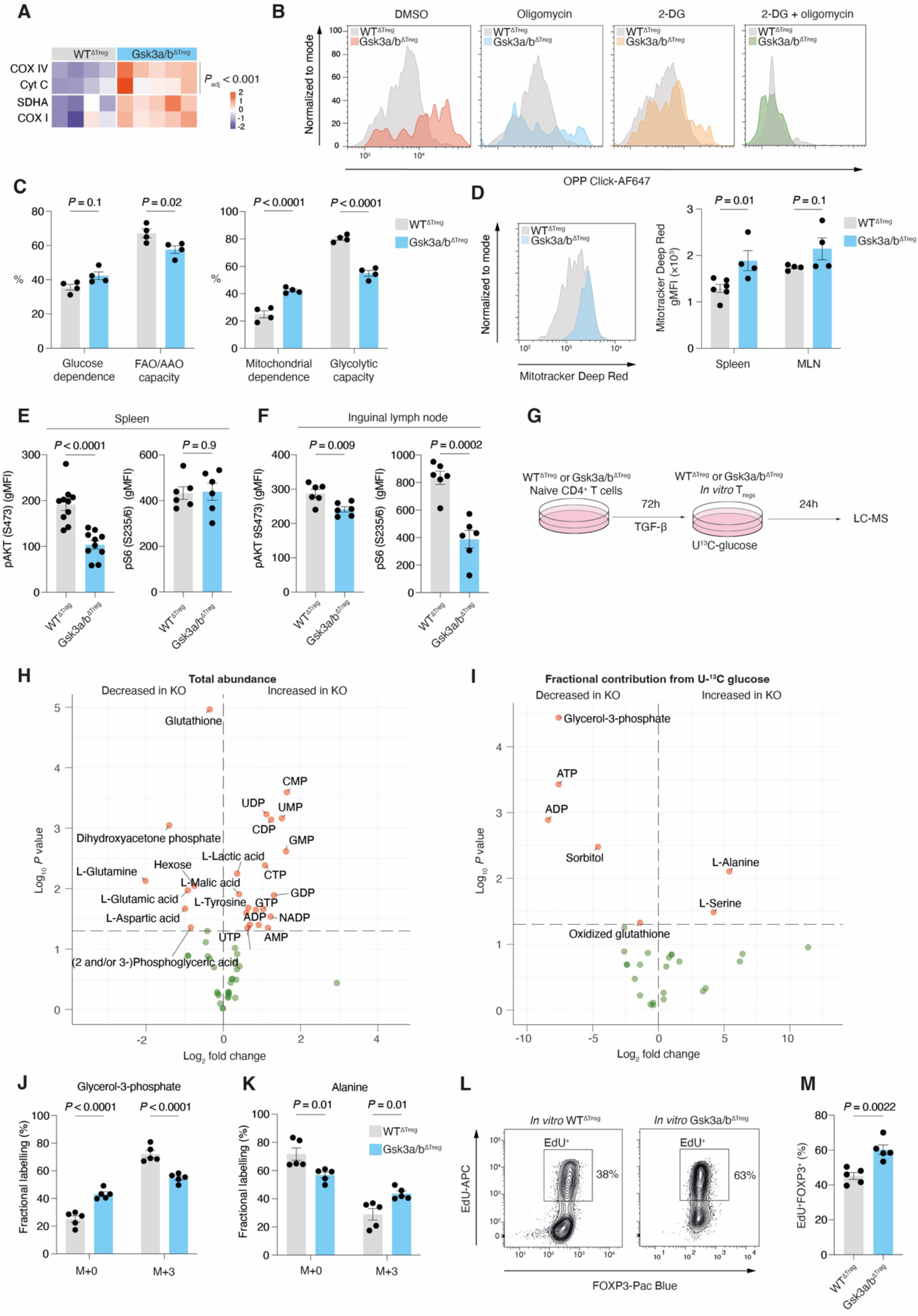
Gsk3a/b deletion rewires T_reg_ metabolism. **A.** Heatmap of gMFI of the indicated mitochondrial proteins, measured by flow cytometry in splenic T_regs_ from WT^ΔTreg^ (n=4) and GSK3a/b^ΔTreg^ mice (n=5). Scale denotes row Z score. Representative of two independent experiments. **B.** Representative histograms of OPP-Click-AF647 fluorescence in WT^ΔTreg^ and Gsk3a/b^ΔTreg^ T_regs_ measured by flow cytometry following treatment with DMSO (vehicle), oligomycin, 2-DG, or 2-DG and oligomycin. **C.** Quantification of glucose dependence and fatty acid/amino acid oxidation dependence, and mitochondrial dependence and glycolytic capacity respectively in WT^ΔTreg^ and Gsk3a/b^ΔTreg^ T_regs_ treated as in B (n=4 per group). Calculations are described in the Methods section. Data pooled from two independent experiments. **D.** Representative histograms and quantification of gMFI of Mitotracker Deep Red fluorescence measured by flow cytometry in WT^ΔTreg^ (n=4-6) and Gsk3a/b^ΔTreg^ T_regs_ (n=4) from spleen and MLN. Data pooled from two independent experiments. **E.** Quantification of gMFI of phospho-AKT (S473) and phospho-S6 (S235/6) in WT^ΔTreg^ (n=6-10) and Gsk3a/b^ΔTreg^ T_regs_ from spleen (n=6-10). Pooled from >3 independent experiments. **F.** Quantification of gMFI of phospho-AKT (S473) and phospho-S6 (S235/6) in WT^ΔTreg^ (n=6-10) and Gsk3a/b^ΔTreg^ T_regs_ from inguinal lymph node (n=6 per group). Pooled from 2 independent experiments. **G.** Schematic of stable isotope labelling experiment. In vitro-T_regs_ were generated, then cultured with U^13^C-glucose for 24 hours before analysis by LC-MS. **H.** Volcano plot of differential total metabolite abundance as in G. WT^ΔTreg^ and Gsk3a/b^ΔTreg^ *in vitro*-T_regs_ were generated from 5 mice (biological replicates) per group. Representative of one independent experiment. **I.** Volcano plot of ^13^C fractional contribution from glucose to indicated metabolites, as in G. WT^ΔTreg^ and Gsk3a/b^ΔTreg^ *in vitro*-T_regs_ were generated from 5 mice (biological replicates) per group. Representative of one independent experiment. **J.** Quantification of glycerol-3-phosphate isotopologues derived from U^13^C-glucose as in G (n=5 biological replicates). Representative of one independent experiment. **K.** Quantification of alanine isotopologues derived from U^13^C-glucose as in G (n=5 biological replicates). Representative of one independent experiment. **L.** Representative flow cytometry plot showing incorporation of EdU into *in vitro*-generated T_regs_ from WT^ΔTreg^ and Gsk3a/b^ΔTreg^ mice. **M.** Quantification of EdU as in **L** (n=5 per group). Data pooled from two independent experiments. Statistical significance was determined by two-way ANOVA with Šidák’s multiple testing correction (A, C-D, J-K) or unpaired two-tailed t test (E-F, M). Data are presented as the mean +/- SEM. Each point represents a single mouse.

Next, to dynamically profile core metabolism in *ex vivo* T_regs_, we used protein translation as a proxy for metabolic rate^39^. Gsk3a/b^ΔTreg^ T_regs_ had a higher baseline rate of incorporation of the puromycin analogue O-propargyl-puromycin (OPP) into nascent proteins, consistent with their more activated state (Fig. 4B). When treated with oligomycin, which inhibits ATP synthase and therefore OxPhos, the reduction in OPP incorporation was more pronounced in Gsk3a/b^ΔTreg^ T_regs_. However, 2-deoxyglucose, which inhibits hexokinase and thus glycolysis, led a larger reduction in OPP uptake in WT^ΔTreg^ T_regs._ These results therefore indicated a higher level of mitochondrial dependence following *Gsk3a/b* deletion, and reduced glycolytic capacity (Fig. 4C). In keeping with this finding, we found increased Mitotracker Deep Red uptake, reflecting higher total mitochondrial mass in Gsk3a/b^ΔTreg^ T_regs_ (Fig. 4D). We then examined AKT phosphorylation, which plays a critical role in T cell metabolic reprogramming and in T_reg_ homeostasis^40,41^. Levels of pAKT (S473) were reduced in both spleen and inguinal lymph node T_regs_, but interestingly phosphorylation of S6, which is a result of mTORC1 activity, along with phospho-mTOR itself, were only downregulated in T_regs_ from inguinal lymph node tissue (Fig. 4E-F, Extended Data Fig. 4A).

We next generated *in vitro*-T_regs_ from Gsk3a/b^ΔTreg^ CD4^+^ T cells, since Gsk3a/b^ΔTreg^ T_regs_ were insufficient in number, and performed stable isotope-resolved liquid chromatography-mass spectrometry (LC-MS) with U-^13^C labelled glucose to profile biochemical intermediates (Fig. 4G). We found that *in vitro* Gsk3a/b^ΔTreg^ T_regs_ had markedly higher levels of predominantly mono- and diphosphate nucleotides (Fig. 4H). Levels of glutathione were lower, as was dihydroxyacetone phosphate (DHAP), glutamine, aspartate, and 2/3-phosphoglyceric acid. Examination of glucose-derived carbon labelling revealed less labelling of glycerol-3-phosphate, ATP, ADP, and sorbitol, and increased ^13^C incorporation into alanine and serine in Gsk3a/b^ΔTreg^ *in vitro* T_regs_ (Fig. 4I-K). Interestingly, although we observed decreased levels of glycolytic intermediates, we did not observe increased glucose-derived ^13^C labelling of TCA intermediates or nucleotides following *Gsk3a/b* deletion, suggesting that alternative carbon sources, such as glutamine may be preferred.

To test whether the increased abundance of nucleotides in Gsk3a/b^ΔTreg^ *in vitro* T_regs_ was reflected in increased DNA synthesis, we pulsed cells with 5-ethynyl-2-deoxyuridine (EdU). We found that EdU incorporation was higher in Gsk3a/b^ΔTreg^ *in vitro* T_regs_ (Fig 4L-M).

Loss of *Gsk3* therefore leads to alteration of T_reg_ metabolism, with increased mitochondrial mass and OxPhos in T_regs_, and *in vitro*, altered nucleotide and glycolytic intermediate abundance.

### Acute deletion of Gsk3a/b leads to tissue-specific activation of T_regs_

Since deletion of *Gsk3* had a profound effect on T_reg_ development, in order to study the role of GSK3 in developmentally intact T_regs_, we next generated mice in which *Gsk3a* and *Gsk3b* could be deleted using tamoxifen-inducible *Foxp3^eGFP^*^-^*^Cre-ERT2^*. To identify cells in which recombination had occurred, we included a *Rosa26*-CAG-LSL-tdTomato allele, which permanently expresses the fluorescent protein tdTomato following Cre-mediated excision of a STOP cassette (hereafter Gsk3a/b^ΔTreg-ERT2^ or WT^ΔTreg-ERT2^ mice)(Fig. 5A). At 14 days following tamoxifen administration, there was highly efficient recombination of the *Rosa26*-CAG-LSL-tdTomato allele, with around 95% of FOXP3^+^ T cells also expressing tdTomato (Fig. 5B). Strikingly there was no difference in the proportions of FOXP3^+^ tdTomato^+^ T_regs_ between Gsk3a/b^ΔTreg-ERT2^ or WT^ΔTreg-ERT2^ mice. We then examined persistence of *Foxp3* expression, by defining ‘ex-T_regs_’ as tdTomato^+^*Foxp3*^eGFP-^, reflecting loss of T_reg_ identity (Fig. 5C). There was a modest increase in ex-T_regs_ in Gsk3a/b^ΔTreg-ERT2^ mice (Fig. 5D-E). We next evaluated expression of key T_reg_ functional proteins. We found no major differences in the levels of CD25, CTLA-4, FOXP3, or NRP-1 in either splenic or inguinal LN Tregs (Extended Data Fig. 5A). However, ICOS, IRF-4, HELIOS, CD39, and β-catenin were all increased in Gsk3a/b^ΔTreg-ERT2^ T_regs_, but only those from the inguinal LN and not the spleen, suggesting a tissue-specific effect in this system (Fig. 5F-J). Notably, β-catenin levels were markedly higher in inguinal LN T_regs_ compared to those from the spleen, in both Gsk3a/b^ΔTreg-ERT2^ and WT^ΔTreg-ERT2^ mice. In keeping with our observations in Gsk3a/b^ΔTreg^ mice, we also found increased mitochondrial mass and membrane potential in Gsk3a/b^ΔTreg-ERT2^ T_regs_, which was also prominent in inguinal LN-derived rather than splenic T_regs_ (Fig. 5K-L).

**Figure 5.**
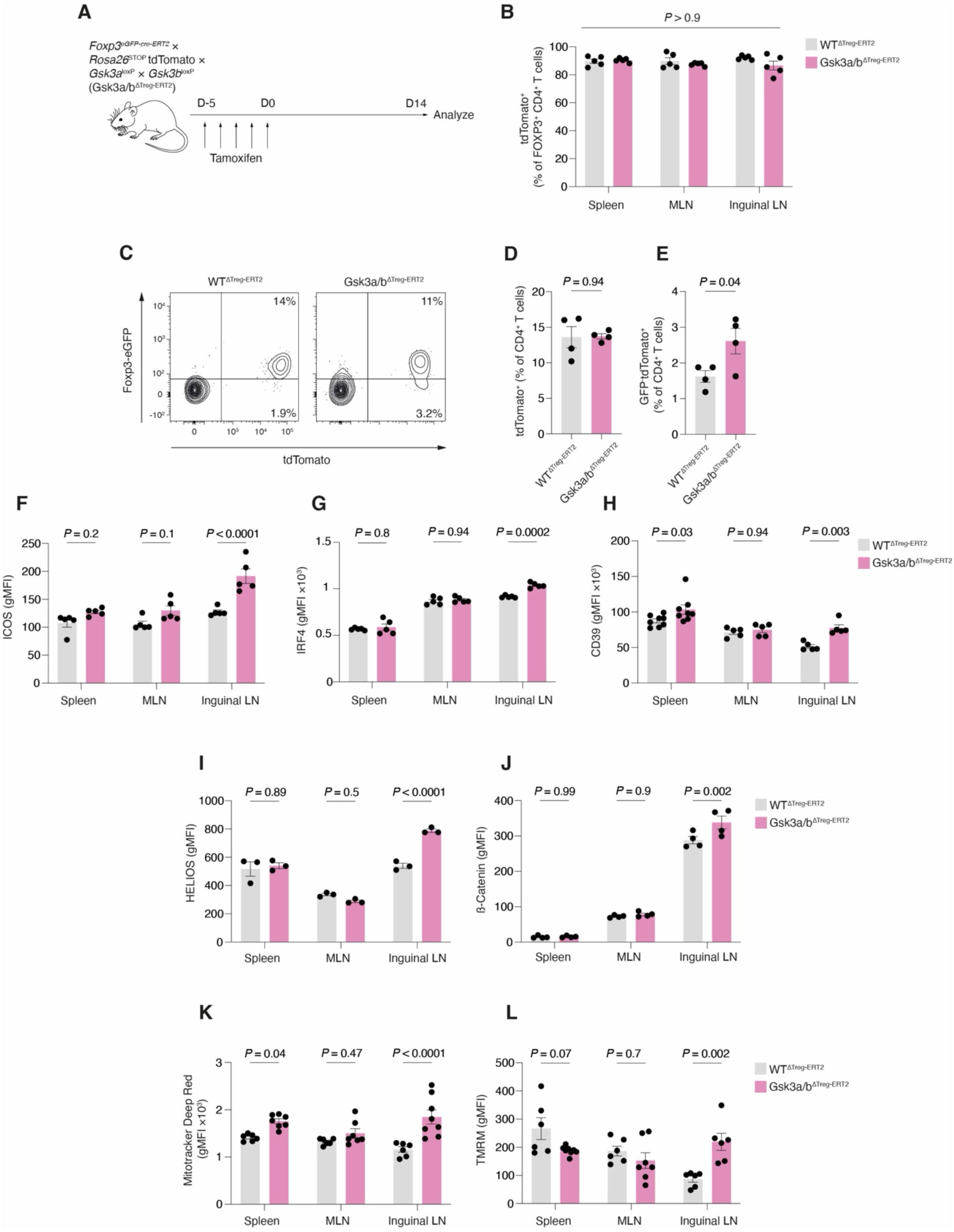
Acute deletion of Gsk3a/b leads to tissue-specific activation of T_regs_. **A.** Schematic of experimental design to induce the deletion of *Gsk3a/b* in regulatory T cells by oral administration of tamoxifen in Gsk3a/b^ΔTreg-ERT2^ mice. **B.** Frequency of tdTomato^+^ cells within the total T_reg_ pool (CD4+FOXP3+) in spleen, MLN, and inguinal lymph nodes of Gsk3a/b^ΔTreg-ERT2^ and WT^ΔTreg-ERT2^ mice at day 14 post-tamoxifen (n=5 mice per group). Representative of two independent experiments. **C.** Representative flow cytometry plot indicating ‘current’ (TCRβ^+^CD4^+^GFP^+^tdTomato^+^) and ‘ex-T_regs_’ (TCRβ^+^CD4^+^GFP^-^tdTomato^+^, based on expression of *Foxp3-eGFP* and tdTomato, in inguinal lymph nodes of of Gsk3a/b^ΔTreg-ERT2^ and WT^ΔTreg-ERT2^ mice at day 14 post-tamoxifen. **D.** Quantification of GFP^+^tdTomato^+^ ‘current’ T_regs_ as in **C**, in inguinal lymph nodes of Gsk3a/b^ΔTreg-ERT2^ and WT^ΔTreg-ERT2^ mice at day 14 post-tamoxifen (n=4 per group). Representative of two independent experiments. **E.** Quantification of GFP^-^tdTomato^+^ ‘ex-T_regs_’ as in **C**, in inguinal lymph nodes of Gsk3a/b^ΔTreg-ERT2^ and WT^ΔTreg-ERT2^ mice at day 14 post-tamoxifen (n=4 per group). Representative of two independent experiments. **F.** Quantification of gMFI of ICOS in TCRβ^+^CD4^+^FOXP3^+^tdTomato^+^ T_regs_ in spleen, MLN, and inguinal lymph nodes of Gsk3a/b^ΔTreg-ERT2^ and WT^ΔTreg-ERT2^ mice at day 14 post-tamoxifen (n=5 per group). Data are pooled from two independent experiments. **G.** Quantification of gMFI of IRF4 in TCRβ^+^CD4^+^FOXP3^+^tdTomato^+^ T_regs_ in spleen, MLN, and inguinal lymph nodes of Gsk3a/b^ΔTreg-ERT2^ and WT^ΔTreg-ERT2^ mice at day 14 post-tamoxifen (n=5 per group). Data are pooled from two independent experiments. **H.** Quantification of gMFI of CD39 in TCRβ^+^CD4^+^FOXP3^+^tdTomato^+^ T_regs_ in spleen, MLN, and inguinal lymph nodes of Gsk3a/b^ΔTreg-ERT2^ and WT^ΔTreg-ERT2^ mice at day 14 post-tamoxifen (n=5-8 per group). Data are pooled from three independent experiments. **I.** Quantification of gMFI of HELIOS in TCRβ^+^CD4^+^FOXP3^+^tdTomato^+^ T_regs_ in spleen, MLN, and inguinal lymph nodes of Gsk3a/b^ΔTreg-ERT2^ and WT^ΔTreg-ERT2^ mice at day 14 post-tamoxifen (n=3 per group). Data are representative of two independent experiments. **J.** Quantification of gMFI of β-catenin in TCRβ^+^CD4^+^FOXP3^+^tdTomato^+^ T_regs_ in spleen, MLN, and inguinal lymph nodes of Gsk3a/b^ΔTreg-ERT2^ and WT^ΔTreg-ERT2^ mice at day 14 post-tamoxifen (n=4 per group). Data are representative of two independent experiments. **K.** Quantification of gMFI of Mitotracker Deep Red in TCRβ^+^CD4^+^FOXP3^+^tdTomato^+^ T_regs_ in spleen, MLN, and inguinal lymph nodes of Gsk3a/b^ΔTreg-ERT2^ and WT^ΔTreg-ERT2^ mice at day 14 post-tamoxifen (n=6-8 per group). Data are pooled from three independent experiments. **L.** Quantification of gMFI of tetramethylrhodamine (TMRM) in TCRβ^+^CD4^+^GFP^+^GITR^hi^ T_regs_ in spleen, MLN, and inguinal lymph nodes of Gsk3a/b^ΔTreg-ERT2^ and WT^ΔTreg-ERT2^ mice (lacking *Rosa26*-CAG-LSL-tdTomato allele) at day 14 post-tamoxifen (n=6-7 per group). Data are pooled from three independent experiments. Statistical significance was determined by two-way ANOVA with Šidák’s multiple testing correction (B, F-L) or unpaired two-tailed t test (D-E). Data are presented as the mean +/- SEM. Each point represents a single mouse.

Acute deletion of Gsk3a/b in T_regs_ therefore does not affect their survival, but induces upregulation of effector proteins predominantly in the inguinal LN, with evidence of loss of a stable identity and metabolic phenotype.

### Gsk3 controls an effector T_reg_ transcriptional program

To explore the effects of acute *Gsk3a/b* deletion on T_reg_ gene expression, we performed single-cell RNA sequencing of sorted *Foxp3*-eGFP^+^tdTomato^+^ T_regs_ isolated from inguinal lymph nodes of Gsk3a/b^ΔTreg-ERT2^ and WT^ΔTreg-ERT2^ mice, at day 14 following completion of tamoxifen administration. This inducible deletion approach allowed detection of transcriptional differences with less potential influence from an inflammatory environment or developmental defects. Following quality control we analysed a total of 16,764 cells from WT^ΔTreg-ERT2^ mice, and 18,746 from Gsk3a/b^ΔTreg-ERT2^ mice. After integration, we defined 8 cell clusters (Fig. 6A-B).

**Figure 6.**
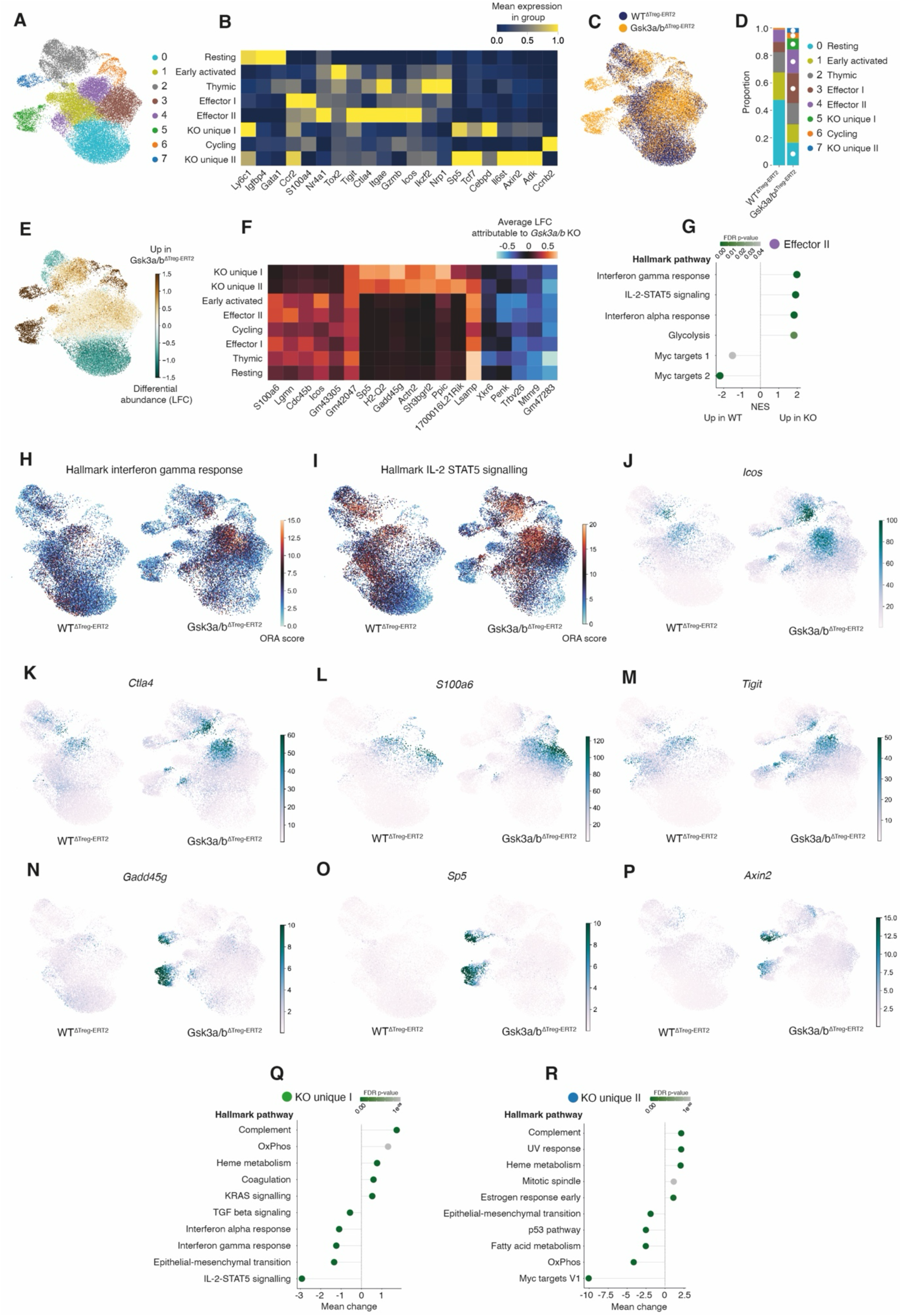
Gsk3 controls an effector T_reg_ transcriptional program. **A.** Uniform manifold approximation and projection (UMAP) plot of integrated dataset of sorted TCRβ^+^CD4^+^GFP^+^tdTomato^+^ T_regs_ from WT^ΔTreg-ERT2^ and Gsk3a/b^ΔTreg-ERT2^ (n=4 per group), at day 14 following tamoxifen course. **B.** Heatmap of marker gene expression for clusters in integrated dataset. **C.** UMAP plot of WT^ΔTreg-ERT2^ and Gsk3a/b^ΔTreg-ERT2^ groups. **D.** Plot of cluster proportions for WT^ΔTreg-ERT2^ and Gsk3a/b^ΔTreg-ERT2^ groups. White dots indicate credible difference (>95% probability). **E.** Differential abundance plot of integrated dataset, coloured by log density. **F.** Heatmap of log fold change differential gene expression across integrated dataset generated by counterfactual analysis of WT^ΔTreg-ERT2^ and Gsk3a/b^ΔTreg-ERT2^ groups, with mapped clusters. **G.** Gene set enrichment analysis (GSEA) of effector II cluster, comparing WT^ΔTreg-ERT2^ and Gsk3a/b^ΔTreg-ERT2^ groups. **H.** Over-representation analysis (ORA) of Hallmark interferon gamma response pathway in individual cells, in WT^ΔTreg-ERT2^ and Gsk3a/b^ΔTreg-ERT2^ groups. **I.** ORA of Hallmark IL-2 STAT-5 signalling pathway in individual cells, in WT^ΔTreg-ERT2^ and Gsk3a/b^ΔTreg-ERT2^ groups. **J.** Normalised expression of *Icos* in WT^ΔTreg-ERT2^ and Gsk3a/b^ΔTreg-ERT2^ groups. **K.** Normalised expression of *Ctla4* in WT^ΔTreg-ERT2^ and Gsk3a/b^ΔTreg-ERT2^ groups. **L.** Normalised expression of *S100a6* in WT^ΔTreg-ERT2^ and Gsk3a/b^ΔTreg-ERT2^ groups. **M.** Normalised expression of *Tigit* in WT^ΔTreg-ERT2^ and Gsk3a/b^ΔTreg-ERT2^ groups. **N.** Normalised expression of *Gadd45g* in WT^ΔTreg-ERT2^ and Gsk3a/b^ΔTreg-ERT2^ groups. **O.** Normalised expression of *Sp5* in WT^ΔTreg-ERT2^ and Gsk3a/b^ΔTreg-ERT2^ groups. **P.** Normalised expression of *Axin2* in WT^ΔTreg-ERT2^ and Gsk3a/b^ΔTreg-ERT2^ groups. **Q.** ORA of Hallmark pathways in KO unique cluster I, relative to all other clusters. **R.** ORA of Hallmark pathways in KO unique cluster II, relative to all other clusters.

Examining the joint dataset, we distinguished a number of shared transcriptional programs. Resting T_regs_ (resting cluster), also represented in topic 1 following latent Dirichlet allocation (LDA)(Extended Data Fig. 6A), expressed *Ly6c1, Igfbp4, and Gata1*^42–44^. The second major program expressed genes associated with T_reg_ activation and effector function (*Ctla4*, *Gzmb*, *Tigit*, *Icos*, *Itgae* and *Prdm1*)(early activated, effector I and effector II clusters; topic 2). The third major program expressed *Ikzf2* (Helios) and *Nrp1*, and represented thymic derived T_regs_ (thymic cluster; topic 3). The relative naivety and activated states of cells in topics 1 and 2 respectively were reflected in their T cell receptor repertoires, with more unique clones in the former compared to the latter (Extended Data Fig. 6B-C).

Examining Gsk3a/b^ΔTreg-ERT2^ T_regs_, we found a marked disturbance of cluster proportions, with an accumulation of effector T_regs_ and a reduction in resting T_regs_(Fig. 6C-D). To explore the effect of *Gsk3a/b* deletion across our complete dataset without annotation, we used a multiresolution variational inference model (MrVI)^45^. Differential abundance analysis again demonstrated a profound shift away from a resting to an activated phenotype, upregulation of a cluster with markers of an active cell cycle, and the emergence of two unique clusters, only found in Gsk3a/b^ΔTreg-ERT2^ T_regs_ (Fig. 6E). We then performed counterfactual differential gene expression measurement, which we cross-referenced with our previous cluster definitions (Fig. 6F). This revealed upregulation of *Icos* and *S100a6* in effector clusters, as well as *Lgmn*, which encodes an asparaginyl endopeptidase important in antigen processing for display on MHC II, and *Cdc45*, required for DNA replication^46,47^. To determine the broader pathways upregulated in the activated clusters, we performed differential gene set enrichment analysis in effector cluster II (Fig. 6G-I). This revealed upregulation of interferon-γ, IL-2-STAT5 signalling, interferon-ɑ, and glycolysis pathways, with downregulation of Myc targets, in Gsk3a/b^ΔTreg-ERT2^ T_regs_ . Projection of key T_reg_ effector gene expression, including *Icos*, *Ctla4*, *S100a6* and *Tigit* onto our data also demonstrated a clear increase in their levels in effector and activated clusters in Gsk3a/b^ΔTreg-ERT2^ T_regs_ (Fig. 6J-M).

There were two unique clusters present only in T_regs_ from Gsk3a/b^ΔTreg-ERT2^ mice. Both clusters expressed elevated *Gadd45g* and *H2-Q2*, as seen in our bulk RNA sequencing analysis of Gsk3a/b^ΔTreg^ T_regs_, and the WNT/β-catenin target genes, *Sp5, Tcf7*, and *Axin2* (Fig. 6N-P). Pathway analysis revealed that despite joint over-representation of complement and heme metabolism gene sets, both clusters had distinct metabolic profiles, with upregulation of an OxPhos gene set in KO unique cluster I, which was downregulated in KO unique cluster II, alongside fatty acid metabolism (Fig. 6Q-R).

Acute deletion of Gsk3 in T_regs_ therefore induces a heterogeneous gene expression program, with upregulation of effector genes, and the emergence of unique clusters expressing genes which may direct a pro-inflammatory phenotype.

### Deletion of Gsk3a/b in T_regs_ leads to enhanced tumour killing and impaired suppressive capacity

Although we had demonstrated increased expression of effector genes and surface proteins in Gsk3a/b^ΔTreg-ERT2^ T_regs_, it remained unclear whether acute rather than constitutive deletion might induce a transient phenotype which enhanced the suppressive capacity of T_regs_. To address this question, we administered tamoxifen to Gsk3a/b^ΔTreg-ERT2^ and WT^ΔTreg-ERT2^ mice, and the next day implanted B16 melanoma cells into their flanks. We then administered further doses of tamoxifen, and followed tumour size over time (Fig. 7A). Gsk3a/b^ΔTreg-ERT2^ mice had dramatically reduced tumour growth (Fig. 7B), with an unconstrained anti-tumour immune response (Fig. 7C-D). There were substantially more CD8^+^ and CD4^+^ tumour-infiltrating lymphocytes (TILs), a high proportion of which produced interferon-γ, but which was not different between genotypes. Within the spleen and inguinal LN however, there was a high proportion of interferon-γ producing CD4^+^ and CD8^+^ T cells in Gsk3a/b^ΔTreg-ERT2^ mice (Fig. 7E-F). In contrast, there was a marked reduction in T_regs_ within the TIL population compared to WT^ΔTreg-ERT2^ mice, but which was not seen in the spleen (Fig. 7G). To confirm the loss of suppressive effect of Gsk3a/b^ΔTreg-ERT2^ T_regs_ seen in an anti-tumoral immune response, we next tested their capacity to suppress T cell proliferation *in vitro*. We generated *in vitro* T_regs_ from naïve Gsk3a/b^ΔTreg-ERT2^ and WT^ΔTreg-ERT2^ CD4^+^ T cells, using 4-hydroxytamoxifen to induce *Gsk3a/b* deletion. We found that in a co-culture assay with wild type responder CD4^+^ T cells, their ability to suppress proliferation was impaired, in common with Gsk3a/b^ΔTreg^ *in vitro* T_regs_. Acute deletion of *Gsk3a/b* therefore impairs T_reg_ function both *in vitro* and *in vivo*.

**Figure 7.**
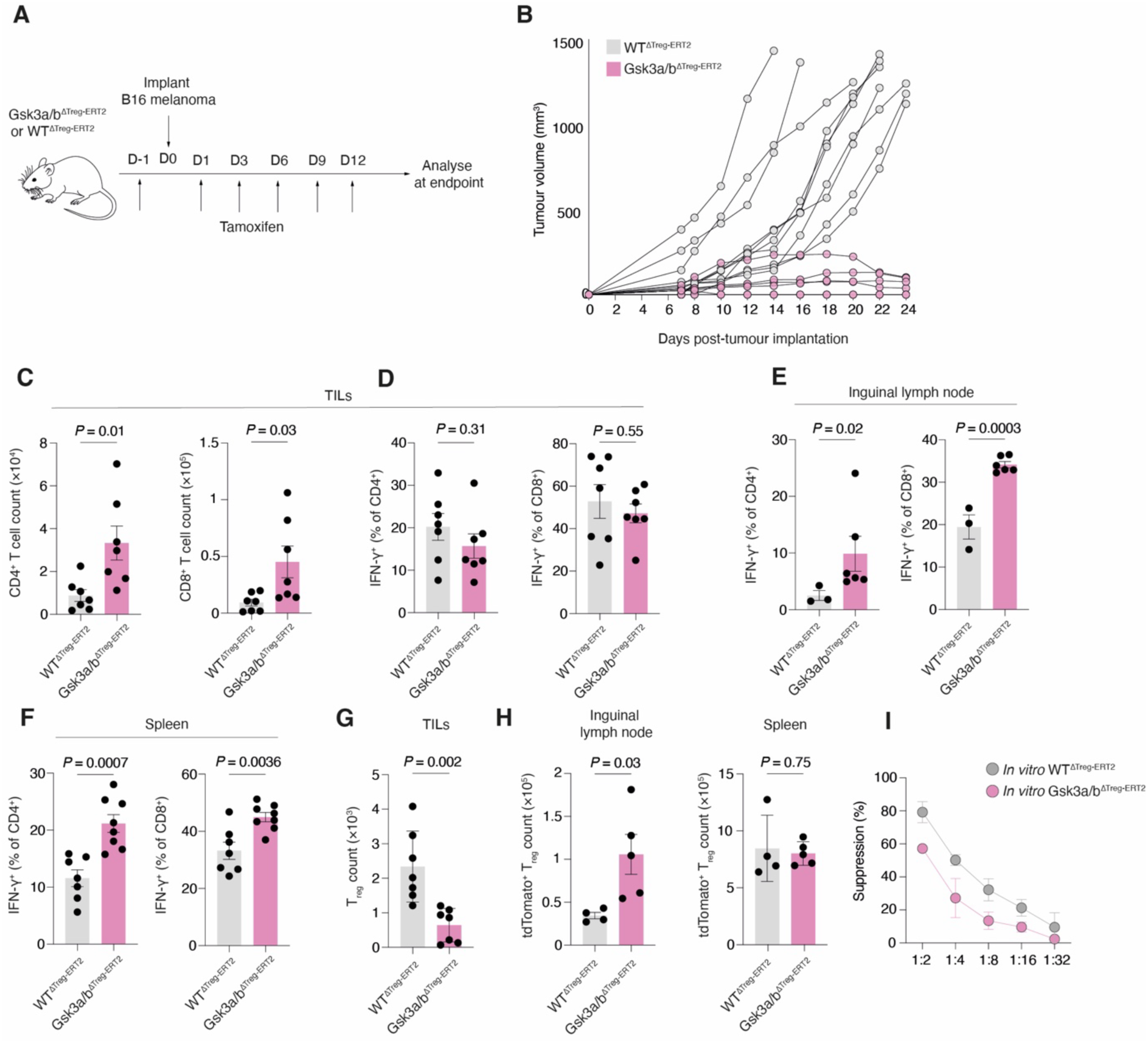
Deletion of Gsk3a/b in T_regs_ leads to enhanced tumour killing and impaired suppressive capacity. **A.** Experimental design for tumour implantation experiments. WT^ΔTreg-ERT2^ and Gsk3a/b^ΔTreg-ERT2^ mice were treated with tamoxifen at day -1, and then B16 melanoma cells implanted, followed by a further 5 doses of tamoxifen, before analysis at endpoint. **B.** Tumour growth curves in WT^ΔTreg-ERT2^ and Gsk3a/b^ΔTreg-ERT2^ mice as in **A** (n=10-14 per group). Each line represents a single mouse. Data pooled from 3 independent experiments. **C.** Quantification of CD4^+^ and CD8^+^ T cell counts in tumour infiltrating lymphocytes (TILs) from WT^ΔTreg-ERT2^ and Gsk3a/b^ΔTreg-ERT2^ mice as in **A** (n=7 per group). Data pooled from two independent experiments. **D.** Quantification of percentage of CD4^+^IFNγ^+^ and CD8^+^IFNγ^+^ T cells in TILs from WT^ΔTreg-ERT2^ and Gsk3a/b^ΔTreg-ERT2^ mice as in **A** (n=7 per group). Data pooled from two independent experiments. **E.** Quantification of percentage of CD4^+^IFNγ^+^ and CD8^+^IFNγ^+^ T cells in inguinal lymph nodes from WT^ΔTreg-ERT2^ and Gsk3a/b^ΔTreg-ERT2^ mice as in **A** (n=3-6 per group). Data pooled from two independent experiments. **F.** Quantification of percentage of CD4^+^IFNγ^+^ and CD8^+^IFNγ^+^ T cells in spleen from WT^ΔTreg-ERT2^ and Gsk3a/b^ΔTreg-ERT2^ mice as in **A** (n=7 per group). Data pooled from two independent experiments. **G.** Quantification of total numbers of FOXP3^+^T_regs_ in TILs from WT^ΔTreg-ERT2^ and Gsk3a/b^ΔTreg-ERT2^ mice as in **A** (n=7 per group). Data pooled from two independent experiments. **H.** Quantification of numbers of tdTomato^+^FOXP3^+^T_regs_ in inguinal lymph node and spleen from WT^ΔTreg-ERT2^ and Gsk3a/b^ΔTreg-ERT2^ mice as in **A** (n=4-5 per group). Data pooled from two independent experiments **I.** Quantification of suppressive capacity of *in vitro*-generated T_regs_ from WT^ΔTreg-ERT2^ and Gsk3a/b^ΔTreg-ERT2^ mice, mixed with wild type responder T cells at the indicated ratio (n=4 per group). Representative of two independent experiments. Statistical significance was determined by unpaired two-tailed t test (C-H). Data are presented as the mean +/- SEM. Each point represents a single mouse.

## Discussion

Our study demonstrates a central role for GSK3 in T_reg_ development and homeostasis, thus identifying it as an essential regulator of immune tolerance. We examined the effect of GSK3 deletion across timescales, using both constitutive and acute deletion models. The former led to profound systemic autoimmunity, associated with tissue-specific and cell-intrinsic numerical defects in T_regs_. Gsk3a/b^ΔTreg^ T_regs_ acquired a transcriptional profile associated with a Th1-like phenotype, with overexpression of *Gadd45g* and *Gzmb*, and metabolic dysregulation. In contrast, acute deletion of *Gsk3a/b* did not affect T_reg_ numbers in the resting state, but caused upregulation of suppression and activation-associated genes, and the emergence of T_reg_ subpopulations expressing β-catenin target genes, which may be precursors of the more dysregulated phenotype seen with constitutive deletion. Despite their potentially suppressive profile, *Gsk3a/b*^-/-^ T_regs_ fail to control the inflammatory response to tumours, or to suppress T_conv_ *in vitro*. These differences highlight divergence between roles for GSK3 in T_reg_ development, compared with their effector function.

An important question is to what extent GSK3 integrates key pathways in T_reg_ biology. One of the most studied targets of GSK3 is the β-catenin degradation axis. Most recent studies show that accumulation of β-catenin, a result of GSK3 inactivation, is detrimental to T_reg_ function. We consistently show upregulation of β-catenin in Gsk3a/b^ΔTreg^ T_regs_, and there are key similarities in differential gene expression when compared with T_regs_ expressing constitutively active β-catenin^21^, including upregulation of *Gadd45g*, *Gzmb*, and *Tmem158*, and downregulation of *Ptger2* and *Cd83*. *Ptger2* encodes the prostaglandin E receptor 2, which has been shown to be upregulated in high-salt environments, such as seen in multiple sclerosis, and which drives β-catenin expression, and may potentially be adaptively downregulated in conditions of β-catenin excess^20^. Deletion of *Ptger2* in T cells leads to enhanced anti-tumour immunity, associated with TCF1 expression^48^, and the low levels of *Ptger2* we observe in Gsk3a/b^ΔTreg^ T_regs_ may contribute to the improved anti-tumoral response to implanted melanoma.

We observed that *Gsk3a/b* deletion had a profound effect on cellular metabolism, with an increase in mitochondrial volume and OxPhos in *ex vivo* cells. Loss of GSK3 in B cells leads to increased oxygen consumption and lactate production, and higher mitochondrial content^49^. In CD8^+^ T cells, GSK3β is found at mitochondrial-endoplasmic reticulum contact sites, and phosphorylates voltage dependent anion channel (VDAC), preventing binding of hexokinase I^50^. One of the most striking findings from our profiling of biochemical intermediates in Gsk3a/b^ΔTreg^ T_regs_ was an accumulation of nucleotides. GSK3 has recently been shown to regulate one-carbon metabolism, a synthetic pathway which serves to transfer one-carbon units with folate as a cofactor, and which is important in nucleotide synthesis^51^. Although we found increased incorporation of ^13^C glucose into serine in Gsk3a/b^ΔTreg^ T_regs_, we did not observe substantial labelling of nucleotides themselves. De novo synthesis of serine is detrimental to T_reg_ function^52^, and this may be a contributory mechanism to their dysfunction in Gsk3a/b^ΔTreg^ mice. An important caveat is that although *in vitro*-generated T_regs_ reproduced key observations in *ex vivo* Gsk3a/b^ΔTreg^ T_regs_, there are potentially significant differences between cell metabolism between these systems.

Acute deletion of *Gsk3a/b* did not lead to a numerical deficiency in T_regs_, but did result in widespread transcriptional alteration, with a profound shift towards an activated phenotype. Gsk3a/b^ΔTreg^ T_regs_ unique clusters expressed β-catenin associated genes, including *Sp5*, *Axin2*, and *Gadd45g*. Despite having transcriptional similarities, these populations had markedly different metabolic profiles, with divergence in their expression of OxPhos and glycolytic gene sets. It is likely that these nascent populations are the precursors of the established GSK3-deficient phenotype we observed in our constitutive deletion model. The underlying drivers of the complex transcriptional changes seen with acute loss of GSK3 remain to be established and are deserving of future investigation.

Previous work has identified an important function for GSK3 in control of anti-tumour immunity. Administration of small molecule GSK3 inhibitors improves tumour killing in vivo, as does deletion of *Gsk3a/b* in all T cells using *Cd4*-Cre^24,28,29^. The effect of these interventions on T_reg_ function has not yet been established, and we speculate that inhibition of their suppressive capacity may represent a significant component of these effects. Moreover, this provides additional rationale to pursue GSK3 inhibition as an adjunct to existing immunotherapy regimes.

## Methods

### Mice

B6.129(Cg)-*Gsk3b^tm2Jrw^*/J (strain 029592), B6.129(Cg)-*Foxp3^tm4(YFP/icre)Ayr^*/J (strain 016959), *Foxp3^tm9(EGFP/cre/ERT2)Ayr^*/J (strain 016961), and B6.Cg-*Gt(ROSA)26Sor^tm9(CAG-tdTomato)Hze^*/J (strain 007909) mice were purchased from the Jackson Laboratory. *Gsk3a*^LoxP^ mice were generated by crossing *Gsk3a*^tm1a(EUCOMM)^ mice with B6.129S4-*^Gt(ROSA)26Sortm1(FLP)Dym^/*RainJ FLP-deleter mice (Jackson Laboratory, strain 009086), to create *Gsk3a*^tm1c(EUCOMM)^ mice^53^.

Male and female mice between the ages of 3 and 7-10 weeks were used equally, except where specified. Mice were bred and maintained under specific pathogen-free conditions at the Kennedy Institute of Rheumatology, University of Oxford. Mice underwent regular checks to ensure they did not carry any pathogenic microorganisms. They were housed in cages that had individual ventilation and were provided with items to stimulate their environment. The temperature was kept between 20 and 24°C, with a humidity level of 45–65%. They were exposed to a 12 h cycle of light and darkness (7:00 to 19:00), with a 30 min period of dawn and dusk. All experiments were performed according to University of Oxford Institutional and UK Home Office regulations under a project license authorized by the UK Home Office (PPL no. PP1971784).

### B16 melanoma model

The B16 melanoma tumour model was performed as previously described^54^. B16-F10 cells were purchased from American Type Culture Collection (ATCC CRL-6475). One day prior to tumour injection (day −1) and on days 1, 3, 6, 9 and 12 days after injection of tumour cells, mice were administered with 4mg of tamoxifen (Sigma) in 100μl corn oil by oral gavage. Two ×10^5^ B16-F10 cells were subcutaneously injected into the flanks of 8-week-old Gsk3a/b^ΔTreg-ERT2^ or WT^ΔTreg-ERT2^ mice. Tumour development and progression was monitored by caliper measurement three times per week. Tumour volume was defined as follows: tumour volume (mm^3^) = L (length) × W (width) × W/2. Mice were culled when their tumour volume reached 1200mm^3^ mm or at 24 days post-implantation, at which point TILs, inguinal lymph nodes and splenic cells were analysed.

### Isolation of immune cells from tumours

To isolate immune cells, tumours were minced and then incubated in RPMI 1640 containing 5% FBS, 1mg/ml collagenase IV (Roche) and 100μg/ml DNase I (Roche) at 37°C for 45 minutes. The digested tumour tissues were then filtered through a 70μm strainer (Falcon) to obtain a single cell suspension. After 2 times washing twice in FACS buffer (PBS containing 0.5% BSA and 2mM EDTA), immune cells were isolated using Ficoll-Paque (Ficoll-Paque Premium 1.084) (Cytiva) density gradient centrifugation. Cells at the interface were removed, and then washed twice with FACS buffer before staining for flow cytometry, and/or stimulation experiments.

### Flow cytometry

Single cell suspensions were prepared from spleen, lymph nodes, thymus and lungs as described previously, with some modifications^55^. Briefly, lungs were finely chopped with scissors before digestion with collagenase type IV (0.5mg/ml) in HBSS with10% FBS at 37°C for 30 minutes. Following digestion, lung tissues were passed through a 70μm cell strainer (Falcon). After two washes in FACS buffer, red blood cell (RBC) lysis was performed in 1ml of ACK RBC lysis buffer (Gibco) for 3 minutes at room temperature (RT), followed by two washes in FACS buffer. Spleens were injected with ice-cold PBS and mashed through a 70μm strainer. Mesenteric lymph nodes, inguinal lymph nodes, and thymus were also mashed through a 70μm strainer. RBCs were depleted by incubating splenocytes with ACK lysis buffer for 3 minutes at RT. For cytokine analysis, cells were stimulated with Cell Stimulation Cocktail (Biolegend) in the presence of brefeldin A and monensin (Biolegend) for 5 hours. Single-cell suspensions were incubated with Fixable Viability Dye eFluor 780 (eBioscience) in PBS (30 minutes at 4°C), followed by Fc Block (TruStain FcX™ (anti-mouse CD16/32) Antibody, Biolegend) (15 mins on ice) and surface antibodies (30 mins on ice) in FACS buffer. For FOXP3 and intracellular staining, cells were fixed and permeabilized using a FOXP3 Staining Kit (eBioscience) following viability and surface antibody staining. For phosphoprotein staining, following viability and surface antibody staining, cells were fixed with FOXP3 fixation and permeabilization buffer for 15 minutes at room temperature, followed by permeabilization with 90% methanol on ice for 30 minutes. Cells were then incubated with Fc block for 15 minutes followed by intracellular staining with phosphoprotein specific antibodies for 30 minutes at room temperature in FACS buffer.

Flow cytometry–based metabolic profiling was performed by the SCENITH method as described^39^, using Click-iT Plus OPP Alexa Fluor 647 Protein Synthesis Assay (Thermo Fisher Scientific). The percentages of mitochondrial dependence, glycolytic capacity, glucose dependence and FAO/AAO (fatty acid oxidation and amino acid oxidation) were measured using the gMFI values of OPP-AF647 from cells treated with 2-deoxyglucose (2-DG), oligomycin, or 2-DG and oligomycin. The formulae are summarized below: Glucose dependence (%): = 100 × [(OPP_No inhib_– OPP_2DG_) / (OPP_No inhib_– OPP_(2DG & oligomycin)_)]. FAO/AAO capacity (%) = 100 - (Glucose Dependence) Mitochondrial dependence (%) = 100 × [(OPP_No inhib_– OPP_oligomycin_) / (OPP_No inhib_– OPP_(2DG & oligomycin)_)] Glycolytic capacity (%) = 100 - (Mitochondrial Dependence).

To assess mitochondrial mass, cells were stained with Fixable Viability Dye at 4 °C followed by surface antibody staining on ice. Then the cells were incubated with MitoTracker Deep Red (Thermo) in RPMI 1640 medium for 30 minutes at 37°C in CO_2_ incubator. For the analysis of Mitochondrial membrane potential, cells were stained with Fixable Viability Dye followed by surface antibody staining. Cells were incubated with TMRM (Thermo Fisher Scientific) at a final concentration of 25 nm for 30 min at 37°C, CO_2_ incubator. Cells were kept warm (at 37°C) until acquired in flow cytometry. Flow cytometry was performed on a BD Fortessa X-20 or LSR II instrument or using a Cytek Aurora (5-laser) spectral flow cytometer. Flow cytometry data were analyzed using FlowJo v10 (BD).

| Antibodies | Manufacturer | Catalogue number |
| --- | --- | --- |
| Brilliant Violet 421 anti-mouse CD19 | Biologend | (BioLegend Cat# 115538, RRID:AB_11203527) |
| PerCP/Cyanine5.5 anti-mouse TCR beta chain | Biologend | (BioLegend Cat# 109228, RRID:AB_1575173) |
| PE/Cyanine7 anti-mouse CD25 | Biologend | (BioLegend Cat# 102016, RRID:AB_312865) |
| Brilliant Violet 605 anti-mouse CD184 (CXCR4) | Biologend | (BioLegend Cat# 146519, RRID:AB_2687359) |
| eFluor 450 anti mouse CD357 (AITR/GITR) | Thermo Fisher Scientific | (Thermo Fisher Scientific Cat# 48-5874-80, RRID:AB_1944395) |
| Brilliant Violet 650 anti-mouse CD69 | Biologend | (BioLegend Cat# 104541, RRID:AB_2616934) |
| Brilliant Violet 510 anti-mouse CD4 | Biologend | (BioLegend Cat# 100553, RRID:AB_2561388) |
| APC anti mouse CD304 (Neuropilin-1) | Thermo Fisher Scientific | (Thermo Fisher Scientific Cat# 17-3041-80, RRID:AB_2573195) |
| PE anti-mouse CD73 | Biologend | (BioLegend Cat# 127205, RRID:AB_1089065) |
| PE/Cyanine7 anti-mouse CD8a | Biologend | (BioLegend Cat# 100721, RRID:AB_312760) |
| Alexa Fluor 700 anti-mouse CD8a | Biologend | (BioLegend Cat# 100730, RRID:AB_493703) |
| PE/Dazzle 594 anti-mouse CD103 | Biologend | (BioLegend Cat# 121430, RRID:AB_2566493) |
| Super Bright 702 anti mouse CD73 | Thermo Fisher Scientific | (Thermo Fisher Scientific Cat# 67-0731-82, RRID:AB_2762796) |
| Brilliant Violet 785 anti-mouse CD279 (PD-1) | Biologend | (BioLegend Cat# 135225, RRID:AB_2563680) |
| Super Bright 702 anti mouse CD25 | Thermo Fisher Scientific | (Thermo Fisher Scientific Cat# 67-0251-80, RRID:AB_2717140) |
| PE-Cyanine7 anti mouse CD39 | Thermo Fisher Scientific | (Thermo Fisher Scientific Cat# 25-0391-82, RRID:AB_1210766) |
| Super Bright 645 anti mouse CD278 (ICOS) | Thermo Fisher Scientific | (Thermo Fisher Scientific Cat# 64-9942-82, RRID:AB_2734943) |
| PerCP-eFluor 710 anti mouse TIGIT | Thermo Fisher Scientific | (Thermo Fisher Scientific Cat# 46-9501-80, RRID:AB_11149128) |
| PE/Cyanine7 anti-mouse CD62L | Biologend | (BioLegend Cat# 104418, RRID:AB_313103) |
| Brilliant Violet 785 anti-mouse TCR beta chain | Biologend | (BioLegend Cat# 109249, RRID:AB_2810347) |
| PE/Dazzle 594 anti-mouse CD73 | Biologend | (BioLegend Cat# 127234, RRID:AB_2800629) |
| Brilliant Violet 650 anti-mouse/human CD44 | Biologend | (BioLegend Cat# 103049, RRID:AB_2562600) |
| Alexa Fluor 647 anti-GL7 Antigen | Biologend | (BioLegend Cat# 144606, RRID:AB_2562185) |
| eFluor 450 anti mouse FOXP3 | Thermo Fisher Scientific | (Thermo Fisher Scientific Cat# 48-5773-82, RRID:AB_1518812) |
| PerCP-eFluor 710 anti mouse FOXP3 | Thermo Fisher Scientific | (Thermo Fisher Scientific Cat# 46-5773-80, RRID:AB_2811809) |
| eFluor 660 anti-mouse beta Catenin | Thermo Fisher Scientific | (Thermo Fisher Scientific Cat# 50-2567-42, RRID:AB_11218086) |
| eFluor 450 anti mouse IRF4 | Thermo Fisher Scientific | (Thermo Fisher Scientific Cat# 48-9858-82, RRID:AB_2574135) |
| eFluor 450 Rat IgG2a kappa Isotype Control | Thermo Fisher Scientific | (Thermo Fisher Scientific Cat# 48-4321-80, RRID:AB_1272002) |
| PE anti-mouse IFN-gamma | Biologend | (BioLegend Cat# 505808, RRID:AB_315402) |
| Rat IgG1 kappa isotype control (eBRG1) | Thermo Fisher Scientific | (Thermo Fisher Scientific Cat# 16-4301-050, RRID:AB_2865969) |
| Alexa Fluor 647 rat IgG1 kappa isotype control (eBRG1) | Thermo Fisher Scientific | (Thermo Fisher Scientific Cat# 51-4301-81, RRID:AB_2802173) |
| PE mouse anti-mTOR (pS2448) | BD Biosciences | (BD Biosciences Cat# 563489, RRID:AB_2736872) |
| Mouse IgG1, k | BD Biosciences | (BD Biosciences Cat# 562292, RRID:AB_11207243) |
| PE anti-mouse phospho-STAT5 (Tyr694) | Thermo Fisher Scientific | (Thermo Fisher Scientific Cat# 12-9010-42, RRID:AB_2572671) |
| PE-Cyanine7 anti-mouse granzyme B | Thermo Fisher Scientific | (Thermo Fisher Scientific Cat# 25-8898-82, RRID:AB_10853339) |
| PerCP-eFluor 710 rat IgG2b kappa isotype control | Thermo Fisher Scientific | (Thermo Fisher Scientific Cat# 46-4031-80, RRID:AB_1834458) |
| PE anti mouse ROR gamma (t) | Thermo Fisher Scientific | (Thermo Fisher Scientific Cat# 12-6981-80, RRID:AB_10805392) |
| PE/Dazzle 594 anti-mouse CD152 (CTLA-4) | Biolegend | (BioLegend Cat# 106318, RRID:AB_2564496) |
| APC anti mouse phospho-S6 (Ser235, Ser236) | Thermo Fisher Scientific | (Thermo Fisher Scientific Cat# 17-9007-42, RRID:AB_2573270) |
| PE-Cyanine7 anti-mouse phospho-mTOR (Ser2448) | Thermo Fisher Scientific | (Thermo Fisher Scientific Cat# 25-9718-41, RRID:AB_2573549) |
| PerCP-eFluor 710 anti mouse Ki-67 | Thermo Fisher Scientific | (Thermo Fisher Scientific Cat# 46-5698-80, RRID:AB_11039489) |
| PE-Vio 615 anti-mouse/human Helios Antibody | Miltenyi Biotec | (Miltenyi Biotec Cat# 130-112-825, RRID:AB_2651983) |
| Alexa Fluor 700 anti-mouse IL-17A | Biolegend | (BioLegend Cat# 506914, RRID:AB_536016) |
| PE-Vio 615 anti mouse CD152 antibody (CTLA-4) | Miltenyi Biotec | (Miltenyi Biotec Cat# 130-116-457, RRID:AB_2727555) |
| eFluor 660 mouse IgG1 kappa isotype control | Thermo Fisher Scientific | (Thermo Fisher Scientific Cat# 50-4714-80, RRID:AB_10598653) |
| PE anti-mouse IFN-gamma | Biolegend | (BioLegend Cat# 505808, RRID:AB_315402) |
| Alexa Fluor 647 anti-mouse TNF-alpha | Biolegend | (BioLegend Cat# 506314, RRID:AB_493330) |
| Brilliant Violet 785 anti-mouse CD19 | Biolegend | (BioLegend Cat# 115543, RRID:AB_11218994) |
| PerCP-eFluor 710 anti mouse Gata-3 | Thermo Fisher Scientific | (Thermo Fisher Scientific Cat# 46-9966-41, RRID:AB_10805385) |
| APC anti mouse T-bet | Thermo Fisher Scientific | (Thermo Fisher Scientific Cat# 17-5825-80, RRID:AB_2744711) |
| PE-Cyanine7 anti-mouse Helios | Thermo Fisher Scientific | (Thermo Fisher Scientific Cat# 25-9883-41, RRID:AB_2637135) |
| PE/Dazzle 594 Armenian hamster IgG isotype control | Biolegend | (BioLegend Cat# 400951, RRID:AB_3096996) |
| PerCP/Cyanine5.5 anti-mouse IL-4 | Biolegend | (BioLegend Cat# 504123, RRID:AB_2561564) |
| PerCP-eFluor 710 Rat IgG2a kappa Isotype Control | Thermo Fisher Scientific | (Thermo Fisher Scientific Cat# 46-4321-80, RRID:AB_1834456) |
| PE-CF594 anti mouse Akt (pS473) | BD Biosciences | (BD Biosciences Cat# 562465, RRID:AB_2737620) |
| APC anti-mouse IFN-gamma | Biolegend | (BioLegend Cat# 505810, RRID:AB_315404) |
| Alexa Fluor 488 anti-GFP | Biolegend | (BioLegend Cat# 338008, RRID:AB_2563288) |
| Alexa Fluor 647 anti-mouse COX IV antibody | Abcam | (Abcam Cat# ab33985, RRID:AB_879754) |
| Alexa Fluor 594 anti-mouse SDHA | Abcam | (Abcam Cat# ab168536, RRID:AB_2857979) |
| Alexa Fluor 647 anti mouse COX I antibody | Abcam | (Abcam Cat#ab198600, clone:1D6E1A8) |
| Alex Fluor 700 anti-mouse cytochrome C | Novus Bio | (Novus Cat# NB100-56503AF700, RRID:AB_3159854) |

### *In vitro* T_reg_ differentiation

Naive CD4^+^ T cells were isolated from mouse spleens by negative selection using MojoSort Mouse CD4 Naïve T Cell Isolation Kits (Biolegend). Plates were coated with 3µg/ml anti-CD3 and 3 µg/ml of anti-CD28 antibody (Biolegend clones 145-2C11 and 37.51) for 2 hours at 37°C, and after washing, 3×10^5^ cells/ml were seeded and cultured in RPMI 1640 (Invitrogen) supplemented with 10% FBS (Gibco), 2 mM L-alanyl-L-glutamine (Glutamax, Gibco), 10 mM HEPES, 1% penicillin/streptomycin, and 50µM β-mercaptoethanol, 5ng/ml of IL-2 IS (improved sequence)(Miltenyi), and 5ng/ml of human TGF-beta1 (Miltenyi) for 6 days. In experiments using Foxp3^eGFP-Cre-ERT2^ mice, cells were isolated and cultured as above, and 0.5µg/ml of 4-hydroxytamoxifen (Sigma) was added at day 3.

### ELISA and cytokine analysis

Mouse IgG1, IgG2a, IgG2b and IgM ELISAs were performed using Thermo ELISA kits, according to manufacturer’s instructions. Serum samples were used in the diluted range of 1:1,000 to 1:10,000, along with appropriate standards. Isotype and IgG subclass detection was done using appropriate secondary HRP-conjugated antibodies with incubation at 37° C for 1 hour. For anti-dsDNA ELISAs, wells Nunc Maxisorp ELISA plates were coated with salmon sperm dsDNA (Sigma D1626) at 100 μg/ml overnight at 4°C. The next day, plates were blocked with 2% BSA in PBS for 1 h at 37°C. Serum at a 1:100 to1:500 dilution in PBS was incubated for 2 h at 37°C, and IgG subclass detection was done using appropriate secondary HRP-conjugated antibodies as described above. After the final washing step, plates were developed with TMB substrate (Sigma, cat: T0440) for 10-30 mins and read on a FLUOstar Omega plate reader. For analysis of cytokines in serum, a LEGENDplex mouse inflammation panel (Biolegend) was used according to manufacturer’s instructions. Serum samples are diluted two fold with assay buffer and run in duplicates. After performing the assay, samples were transferred into cluster tubes and acquired by flow cytometry on the same day. Data were analyzed using LEGENDplex data analysis software (Biolegend).

### Immunoblotting

2 × 10^6^ *in vitro*-generated regulatory T cells were washed twice in ice-cold PBS, resuspended in 100μl of RIPA lysis buffer with cOmplete™ EDTA-free Protease Inhibitor and PhosSTOP (Roche) and incubated at 4°C for 15 minutes with frequent vortexing. After centrifugation at 15,000×g for 15 minutes, the supernatant was removed and protein concentration was measured using a BCA kit (BioRad). Protein samples were denatured using Laemmli sample buffer (BioRad) with 10% β-mercaptoethanol (Sigma) and heated to 95°C for 5 minutes. 20μg of protein was run on a MiniProtean TGX gel (BioRad) in SDS buffer. Protein was transferred to a PVDF membrane (Millipore) using a Trans-Blot Turbo transfer system (BioRad). After transfer, the membrane was blocked with 5% non-fat milk in tris buffered saline (TBS) at room temperature for 1 h. The membrane was then probed with rabbit anti-mouse vinculin antibody (Cell Signalling Technology, Cat #13901S) and rabbit anti-mouse GSK3α/β antibody (Cell Signalling Technology, Cat #5676) overnight at 4°C. After washing, the membrane was incubated with anti-rabbit IgG, HRP-linked Antibody (Cell Signalling Technology, Cat#7074) in 1:20,000,2.5% milk TBS-T with 0.01% SDS for 90 minutes at room temperature. After washing five times in TBS-T, membranes were developed using ECL western blot detection reagents (Pierce), and signal was detected by radiographic film (Cytiva).

### *In vitro* T cell suppression assay

Responder cells were prepared by isolating naive CD4+ T cells as previously described, which were labelled with CellTrace Violet (CTV) using the CTV Cell Proliferation Kit (Thermo) according to manufacturer’s instructions. *In vitro*-differentiated T_regs_ and responder CD4^+^ T cells were co-cultured in ‘U’ bottom 96 well plates at the indicated ratios in the presence of washed Dynabeads Mouse T cell CD3/CD28 Activator Beads (Thermo), at a bead-to-cell ratio of 1:1 for 3 days. As a positive control, responder cells were stimulated without T_regs_. Percentage suppression was calculated using the following formula: 100 -[100 × (percentage of proliferating cells with T_regs_ present)/(percentage of proliferating cells without T_regs_ present)].

### Metabolomics

*In vitro* T_regs_ were differentiated as described above, for 4 days. On day 4, cells were washed twice in PBS and resuspended in glucose-free RPMI containing 10% dialyzed FBS, 2mM L-glutamine, 10mM HEPES, 1% penicillin/streptomycin, 50µM β-mercaptoethanol, 5ng/ml of IL-2,5ng/ml of TGF beta1 and uniformly labelled [^13^C]-Glucose (Cambridge Isotope Laboratories) at 10μM. After 24hrs, cells were washed twice in ice-cold PBS. Metabolites were then extracted with ice-cold 80% LC-MS grade-methanol (Sigma), and then vacuum-dried using a Speedvac.

Samples were loaded into a Dionex UltiMate 3000 LC System (Thermo Scientific Bremen, Germany) equipped with a C-18 column (Acquity UPLC-HSS T3 1. 8 µm; 2.1 x 150 mm, Waters) coupled to a QExactive Orbitrap mass spectrometer (Thermo Scientific) operating in negative ion mode. A step gradient was carried out using solvent A (10mM TBA and 15mM acetic acid) and solvent B (100% methanol). The gradient started with 5% of solvent B and 95% solvent A and remained at 5% B until 2 min post injection. A linear gradient to 37% B was carried out until 7 min and increased to 41% until 14 min. Between 14 and 26 minutes the gradient increased to 95% of B and remained at 95% B for 4 minutes. At 30 min the gradient returned to 5% B. The chromatography was stopped at 40 min. The flow was kept constant at 0.25 mL/min and the column was placed at 40°C throughout the analysis. The MS operated in full scan mode (m/z range: [70.0000-1050.0000]) using a spray voltage of 4.80kV,capillary temperature of 300°C,sheath gas at 40.0, auxiliary gas at 10.0. The AGC target was set at 3.0E+006 using a resolution of 140000, with a maximum IT fill time of 512 ms. Data collection was performed using the Xcalibur software (Thermo Scientific). The data analyses were performed by integrating the peak areas (El-Maven– Polly-Elucidata).

Differential abundance of metabolites was determined with linear modelling using the R packages *limma* and *voom*.

### Bulk RNA sequencing

*Foxp3*-YFP^+^ cells were FACS sorted from the spleens of 3-week-old Gsk3a/b^ΔTreg^ and WT^ΔTreg^ mice following CD4^+^ T cell enrichment (MojoSort Mouse CD4^+^ T cell Isolation Kit, Biolegend) by negative selection. Around 10,000 cells were sorted per mouse. RNA was isolated using an RNeasy Plus Micro Kit (Qiagen). RNA was quantified using RiboGreen (Invitrogen) on a FLUOstar OPTIMA plate reader (BMG Labtech) and the size profile and integrity analysed on a 2200 or 4200 TapeStation (Agilent, RNA ScreenTape). RIN estimates were between 9.3 and 9.7. Input material was normalised to 100 ng prior to library preparation. Polyadenylated transcript enrichment and strand specific library preparation was completed using NEBNext Ultra II mRNA kit (NEB) following manufacturer’s instructions. Libraries were amplified (17 cycles) on a Tetrad (Bio-Rad) using in-house unique dual indexing primers. Individual libraries were normalised using Qubit, and the size profile was analysed on a 2200 or 4200 TapeStation. Individual libraries were normalised and pooled together accordingly. The pooled library was diluted to ∼10nM for storage. The 10nM library was denatured and further diluted prior to loading on the sequencer. Paired end sequencing was performed using a NovaSeq6000 platform at the Centre for Human Genetics, University of Oxford (Illumina, NovaSeq 6000 S2/S4 reagent kit, 300 cycles). Post-sequencing quality control and adapter trimming was performed with *fastp*, transcripts were counted using *Salmon*, and differential gene expression and pathway analysis performed using the R package *DESeq2* and the Python package *decoupler*. Genes or pathways were considered significantly different if false discovery rate adjusted *P* value was <0.05.

### Single cell RNA sequencing

Following flow sorting, viable tdTomato^+^ cells from individual mice were labelled with TotalSeq^TM^-C anti-mouse Hashtag oligonucleotide-conjugated antibodies (Biolegend) and combined into two pools, each containing equal numbers of Gsk3a/b^ΔTreg-ERT2^ and WT^ΔTreg-ERT2^ biological replicates. Each pool was then loaded onto a 10X Genomics Chromium Controller (Chip K). Gene expression and feature barcode libraries were prepared using the 10X Genomics Single Cell 5′ Reagent Kit v2 according to the manufacturer’s user guide (CG000330 Rev B). The final libraries were diluted to approximately 10nM for storage. The 10nM library was denatured and further diluted before loading on a NovaSeq 6000 sequencing platform (v.1.5 chemistry, 28/98 bp paired-end, Illumina) at Novogene (Cambridge, UK).

Filtered matrices were generated using Cellranger 7.1 and pre-processed with *Scanpy*. Individual samples were demultiplexed and doublets removed using *Solo*^56^. Cells with fewer than 300 genes, or genes found in fewer than 10 cells were removed, as were those with more than 7% mitochondrial genes. Sequencing batches were integrated using *scVI tools* (n_hidden = 128, n_latent=10, and n_layers=1)^57^, and the resulting latent representation used for clustering and visualisation. Cluster marker genes were identified using the differential gene expression function of *scVI tools* followed by manual annotation based on canonical markers and reference to the literature. Differential cluster abundance was determined using *scCODA*^58^. Multiresolution variational inference was implemented using *MrVI*^45^. Pathway analysis comparing genotypes was performed using *decoupler* (1.6), following pseudobulking of the examined cluster, and over-representation analysis was used to determine pathway scores for single cells, again using *decoupler*. TCR sequences were assembled from 10X gene expression sequencing outputs using TRUST4, and analysed using *scRepertoire*^59^.

### Statistical analysis

The use of the statistical tests is indicated in the respective figure legends, with the error bars indicating the mean ± S.E.M. P values ≤ 0.05 were considered to indicate significance. Analyses were performed with GraphPad Prism v10 or R 4.1. No statistical methods were used to pre-determine sample sizes, but our sample sizes are similar to those reported in previous publications. The distribution of data was determined using normality testing to determine appropriate statistical methodology, or otherwise assumed to be normally distributed. For *in vivo* experiments we matched the sex and age of the mice in experimental batches, but other modes of randomization was not performed. Data collection and analysis were not performed blind to the conditions of the experiments in most of the experiments. Mice with complete absence of germinal centers as well as lack of Alum spots after immunization were considered as failed intraperitoneal immunization and therefore excluded from analysis.

## Data availability

RNA sequencing data has been deposited to GEO under accession GSE277605

## Code availability

Code used in analyzing single cell RNA sequencing will be uploaded to: https://github.com/alexclarke7

## Acknowledgements

We thank Jonathan Webber for flow sorting, and we are also grateful to the Kennedy Institute BSU staff for their support. We thank VIB Metabolomics Centre for performing metabolomics experiments.

Funding for this work was provided by the Wellcome Trust (211072/Z/18/Z) and Cancer Research UK/Versus Arthritis (C70663/A29547) to A.J.C. Flow cytometry and microscopy facilities were supported by the Kennedy Trust for Rheumatology Research through the Cell Dynamics Platform. The computational aspects of this research were supported by the Wellcome Trust (Core Award Grant Number 203141/Z/16/Z) and the NIHR Oxford BRC. The views expressed are those of the author(s) and not necessarily those of the NHS, the NIHR or the Department of Health. The image of a laboratory mouse used was created by Gwilz and distributed under a CC BY-SA 4.0 license. For the purpose of open access, the author has applied a CC BY-ND public copyright license to any Author Accepted Manuscript version arising from this submission.

## Author Contributions

M.K. and A.C. conceived and designed the study. M.K. performed the majority of the experiments. R.M., H.A., M.A. and M.B. performed experiments. A.C. and I.R. analyzed data. A.C. supervised the study.

---

**Extended Data Figure 1.**
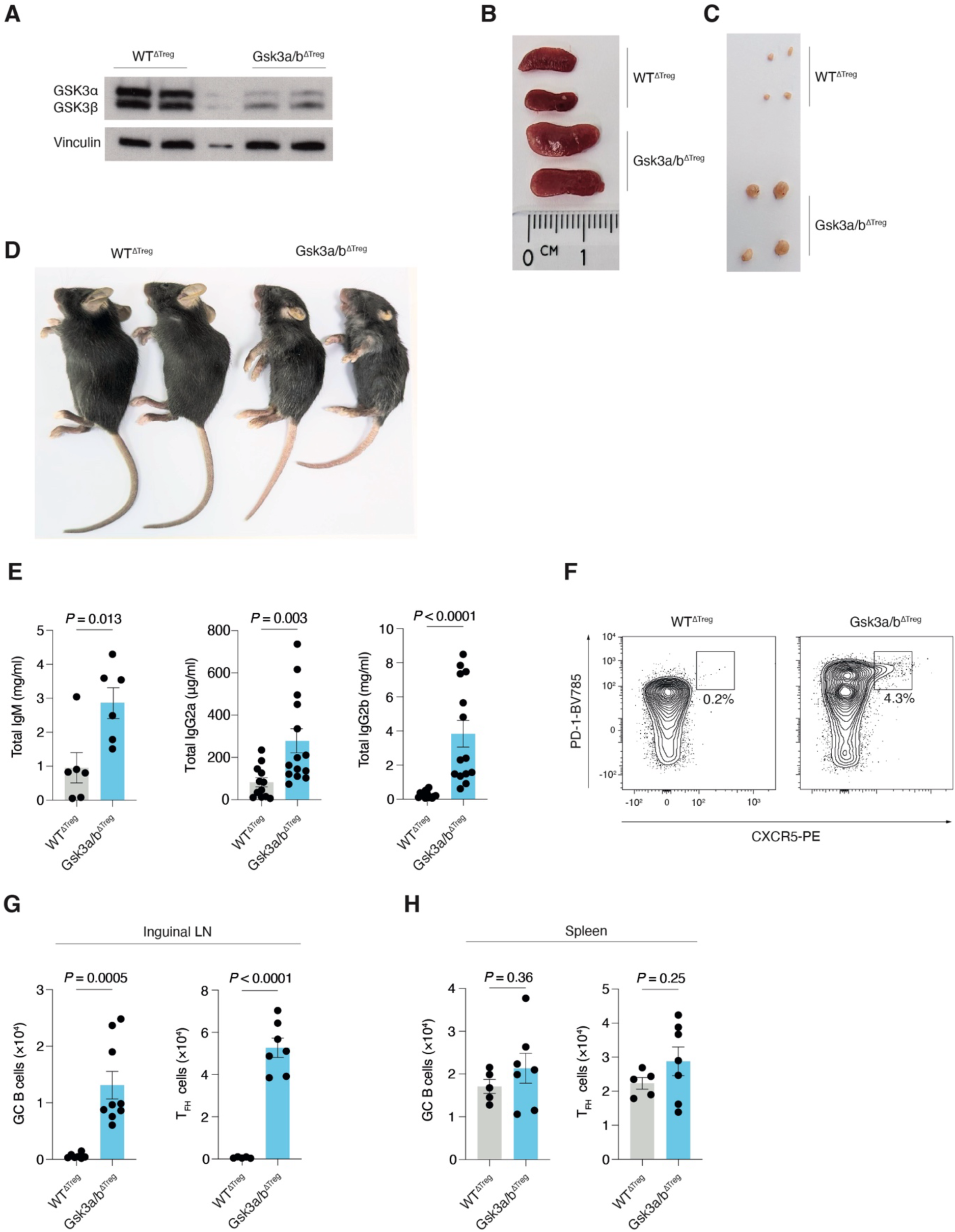
**A.** Immunoblot of GSK3ɑ/β from *in vitro*-generated T_regs_ from WT^ΔTreg^ and Gsk3a/b^ΔTreg^ mice. Representative of two independent experiments. **B.** Representative photographs of the spleens of WT^ΔTreg^ and Gsk3a/b^ΔTreg^ mice at 3 weeks of age. **C.** Representative photographs of the inguinal lymph nodes of WT^ΔTreg^ and Gsk3a/b^ΔTreg^ mice at 3 weeks of age. **D.** Representative photographs of WT^ΔTreg^ and Gsk3a/b^ΔTreg^ mice at 3 weeks of age. **E.** ELISA quantification of total serum IgM, IgG2a, and IgG2b in WT^ΔTreg^ and Gsk3a/b^ΔTreg^ mice by ELISA (n=6-14 per group).Pooled from >5 independent experiments. **F.** Representative flow cytometry gating strategy for T_FH_ cells (CD4^+^PD-1^hi^CXCR5^hi^). **G.** Quantification of GC B cell and T_FH_ cell counts in inguinal lymph nodes of WT^ΔTreg^ and Gsk3a/b^ΔTreg^ mice at 3 weeks of age (n=5-9 per group). Data pooled from two independent experiments. **H.** Quantification of GC B cell and T_FH_ cell counts in spleens of WT^ΔTreg^ and Gsk3a/b^ΔTreg^ mice at 3 weeks of age (n=5-7 per group). Data pooled from two independent experiments. Statistical significance was determined by unpaired two-tailed t test (E,G-H). Data are presented as the mean +/- SEM. Each point represents a single mouse.

**Extended Data Figure 2.**
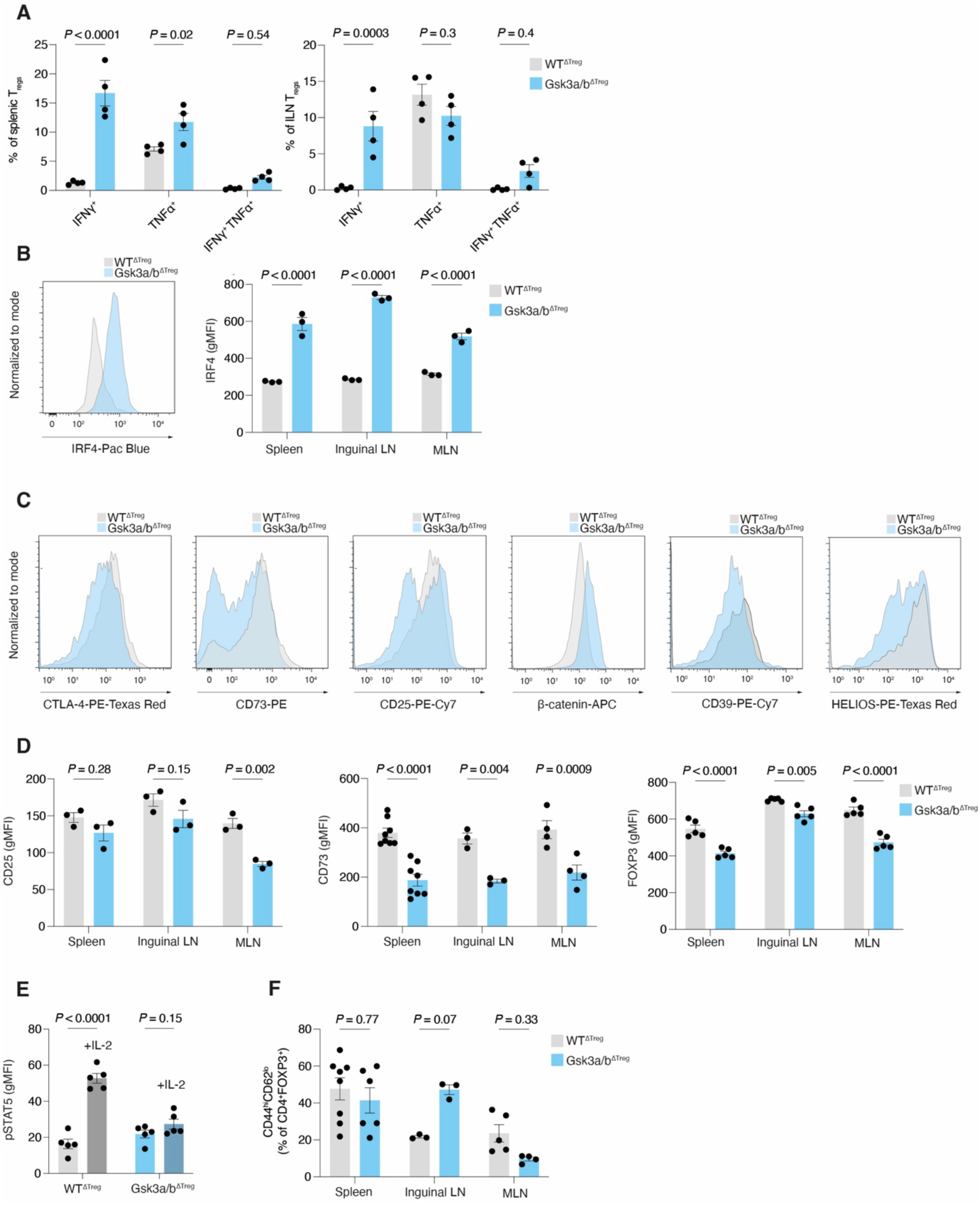
**A.** Quantification of frequency of IFNγ^+^,TNFα^+^, or IFNγ^+^TNFα^+^ T_regs_ in spleen and inguinal lymph nodes of WT^ΔTreg^ and Gsk3a/b^ΔTreg^ mice (n=4 per group). Pooled from two independent experiments. **B.** Representative histogram of IRF4 fluorescence, and quantification of IRF4 gMFI in T_regs_ from WT^ΔTreg^ and Gsk3a/b^ΔTreg^ mice in spleen, inguinal lymph nodes, and MLN (n=3 per group). Representative of two independent experiments. **C.** Representative histograms of CTLA-4, CD73, CD25, β-catenin, CD39, and HELIOS fluorescence in T_regs_ from WT^ΔTreg^ and Gsk3a/b^ΔTreg^ mice in spleen. **D.** Quantification of CD25, CD73, and FOXP3 gMFI in T_regs_ from WT^ΔTreg^ and Gsk3a/b^ΔTreg^ mice in spleen, inguinal lymph nodes, and MLN (n=3-8 per group). Data pooled from 2 independent experiments. **E.** Quantification of phospho-STAT5 in splenic T_regs_ from WT^ΔTreg^ and Gsk3a/b^ΔTreg^ mice stimulated with IL-2 (n=5 per group). Representative of two independent experiments. **F.** Quantification of CD44^hi^ CD62L^lo^ T_regs_ from spleen, inguinal lymph node, and MLN in WT^ΔTreg^ and Gsk3a/b^ΔTreg^ mice (n=3-7 per group). Representative of, or pooled from >2 independent experiments. Statistical significance was determined by two-way ANOVA with Šidák’s multiple testing correction (A-B, D-F). Data are presented as the mean +/- SEM. Each point represents a single mouse.

**Extended Data Figure 3.**
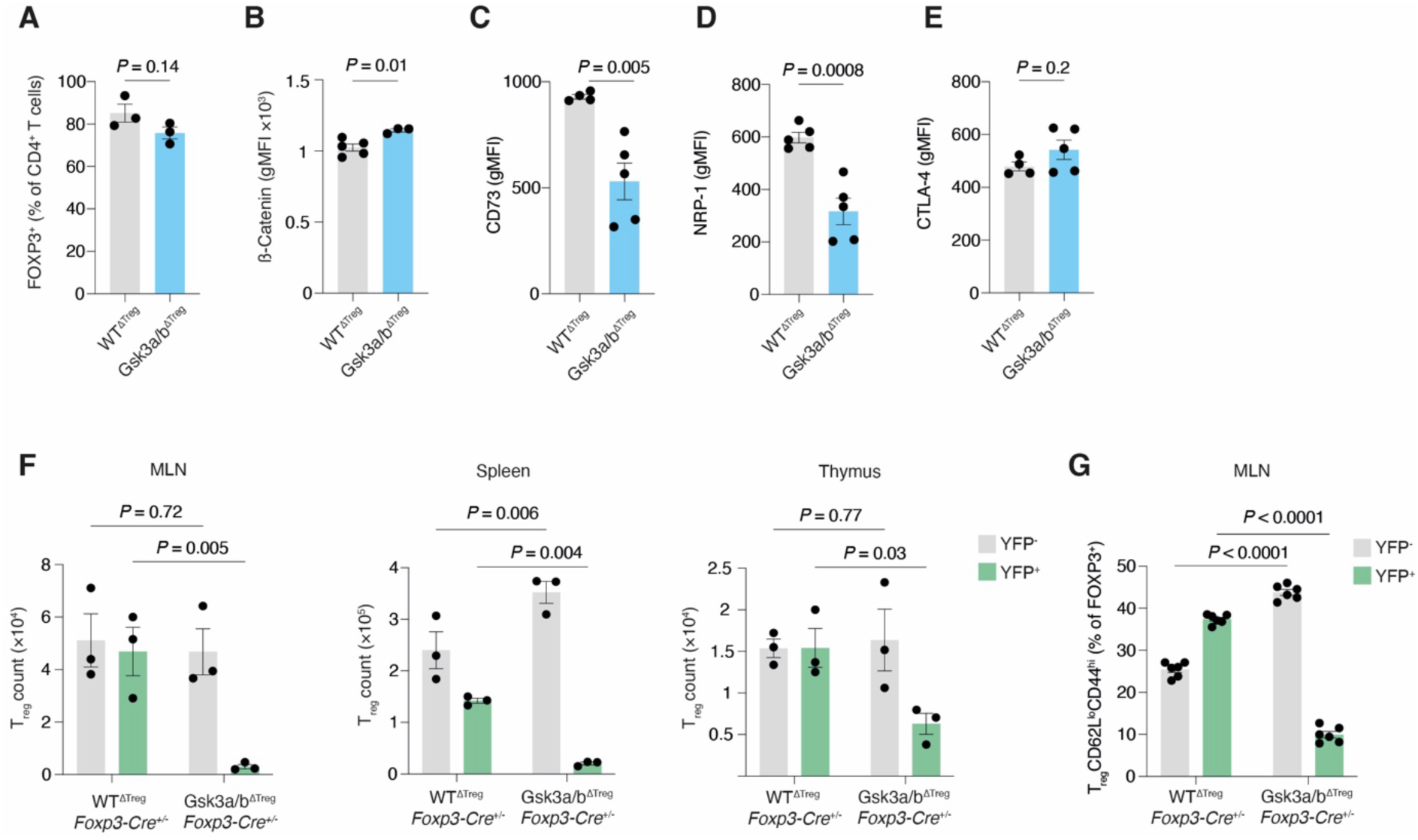
**A.** Quantification of FOXP3^+^ T cells in *in vitro*-generated T_regs_ from WT^ΔTreg^ or Gsk3a/b^ΔTreg^ mice (n=3 per group). Representative of two independent experiments. **B.** Quantification of β-catenin in *in vitro*-generated FOXP3^+^ T_regs_ from WT^ΔTreg^ or Gsk3a/b^ΔTreg^ mice (n=3 per group). Representative of two independent experiments. **C.** Quantification of CD73 in *in vitro*-generated FOXP3^+^ T_regs_ from WT^ΔTreg^ or Gsk3a/b^ΔTreg^ mice (n=3 per group). Representative of two independent experiments. **D.** Quantification of NRP-1 in *in vitro*-generated FOXP3^+^ T_regs_ from WT^ΔTreg^ or Gsk3a/b^ΔTreg^ mice (n=5 per group). Pooled from two independent experiments. **E.** Quantification of CTLA-4 in *in vitro*-generated FOXP3^+^ T_regs_ from WT^ΔTreg^ or Gsk3a/b^ΔTreg^ mice (n=5 per group). Pooled from two independent experiments. **F.** Quantification of absolute counts of YFP^+^ or YFP^-^ FOXP3^+^ T_regs_ from female WT^ΔTreg^ *Foxp3*^YFP^-Cre^+/-^ or Gsk3a/b^ΔTreg^ *Foxp3*^YFP^-Cre^+/-^ mice in MLN, spleen or thymus (n=5 per group). Representative of two independent experiments. **G.** Quantification of frequency of CD44^hi^CD62L^lo^ T_regs_ in female WT^ΔTreg^ *Foxp3*^YFP^-Cre^+/-^ or Gsk3a/b^ΔTreg^ *Foxp3*^YFP^-Cre^+/-^ mice in MLN (n=6 per group). Data pooled from two independent experiments. Statistical significance was determined by unpaired two-tailed t test (A-E) or two-way ANOVA with Šidák’s multiple testing correction (F). Data are presented as the mean +/- SEM. Each point represents a single mouse.

**Extended Data Figure 4.**
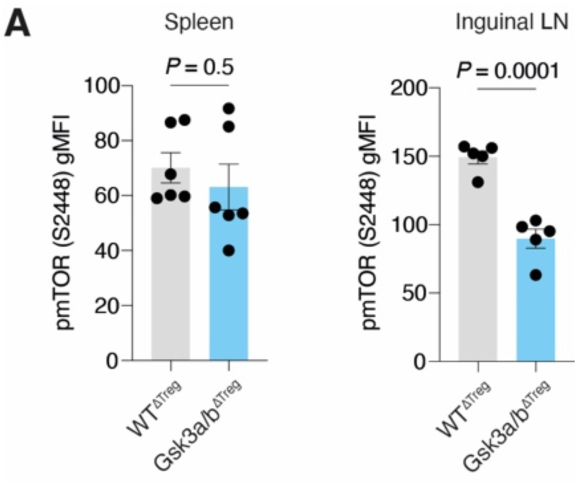
**A.** Quantification of gMFI of phospho-mTOR (S2448) in T_regs_ from spleen and inguinal lymph nodes of WT^ΔTreg^ or Gsk3a/b^ΔTreg^ mice. Data shown are mean± SEM. Data are pooled from two independent experiments (n=5-6 mice). Statistical significance was determined by unpaired two-tailed t test. Data are presented as the mean +/- SEM. Each point represents a single mouse.

**Extended Data Figure 5.**
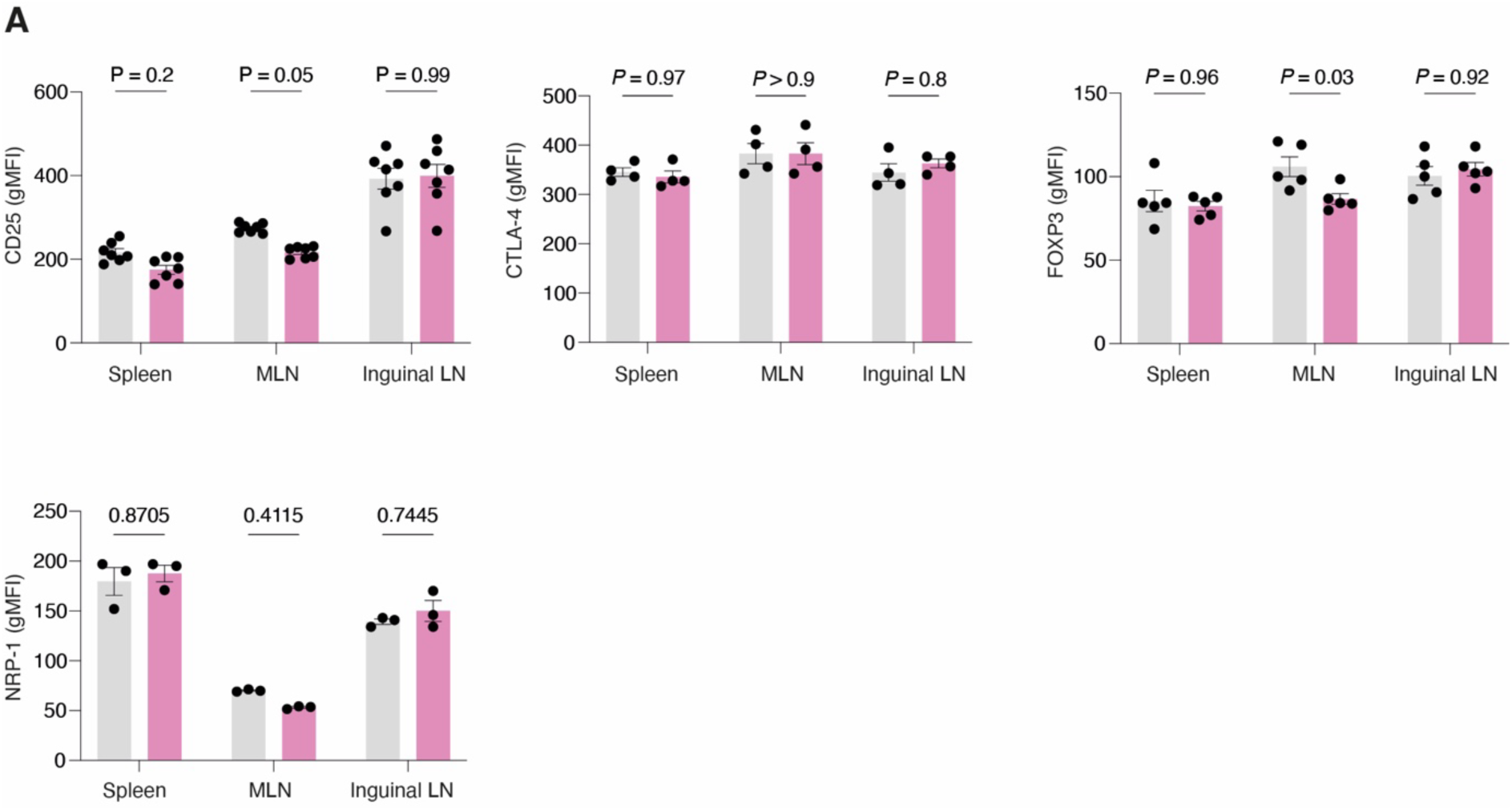
**A.** Quantification of gMFI of CD25, CTLA-4, FOXP3, and NRP-1 in T_regs_ from in spleen, MLN, and inguinal lymph nodes of Gsk3a/b^ΔTreg-ERT2^ and WT^ΔTreg-ERT2^ mice at day 14 post-tamoxifen (n=3-7 per group). Data are pooled from two independent experiments. Statistical significance was determined by two-way ANOVA with Šidák’s multiple testing correction. Data are presented as the mean +/- SEM. Each point represents a single mouse.

**Extended Data Figure 6.**
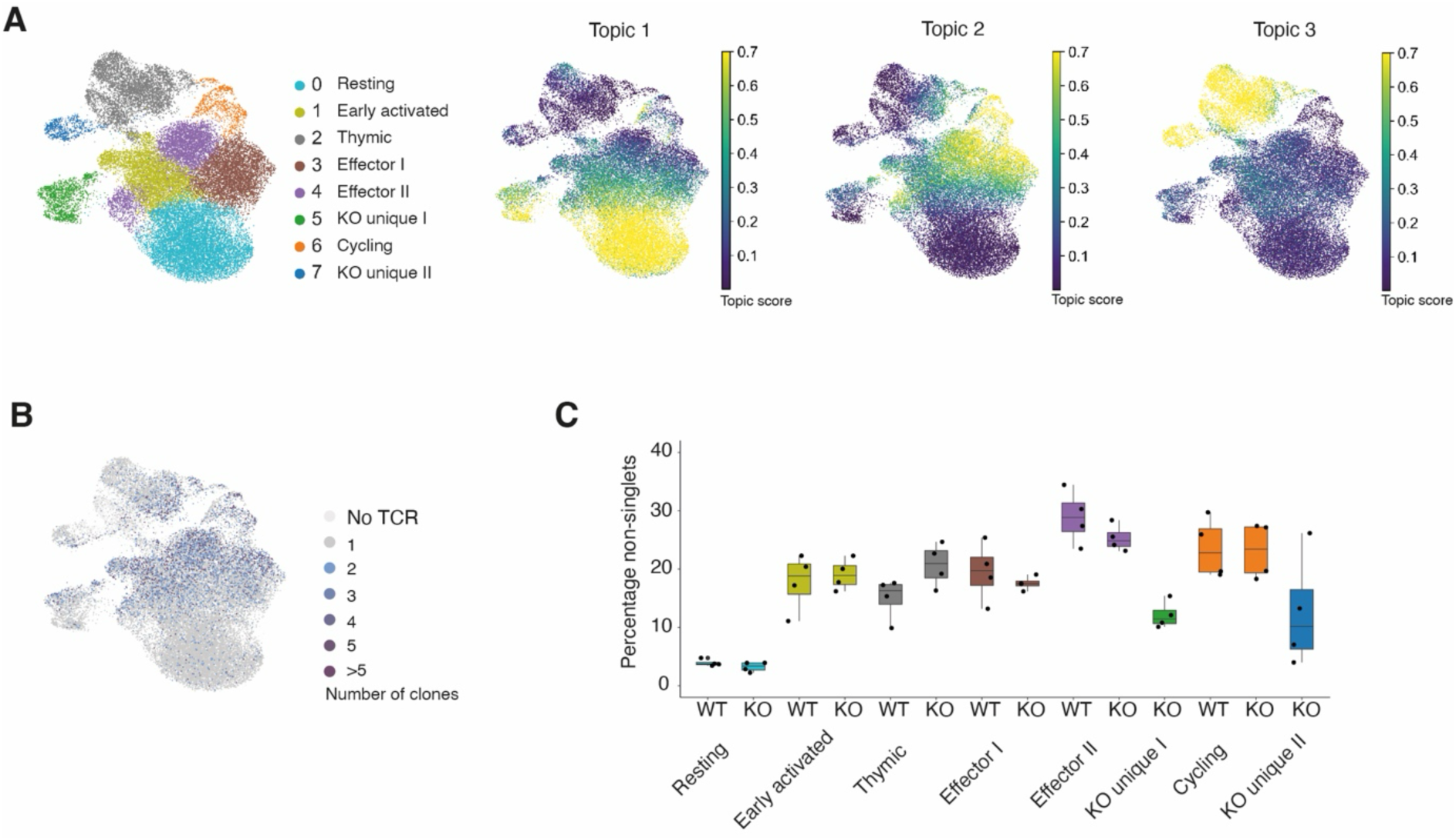
**A.** UMAP projections of T_reg_ clusters, and feature plot of topic scores for LDA topics 1-3 in WT^ΔTreg-ERT2^ and Gsk3a/b^ΔTreg-ERT2^ mice as in Figure 6. **B.** Feature plot of TCR clonality. **C.** Quantification of percentage of non-singlets (and therefore clonal) TCRs, by cluster in WT^ΔTreg-ERT2^ and Gsk3a/b^ΔTreg-ERT2^ mice.

